# Automated CUT&Tag profiling of chromatin heterogeneity in mixed-lineage leukemia

**DOI:** 10.1101/2020.10.06.328948

**Authors:** Derek H. Janssens, Michael P. Meers, Steven J. Wu, Ekaterina Babaeva, Soheil Meshinchi, Jay F. Sarthy, Kami Ahmad, Steven Henikoff

## Abstract

Acute myeloid and lymphoid leukemias often harbor chromosomal translocations involving the *Mixed Lineage Leukemia-1* gene, encoding the KMT2A lysine methyltransferase. The most common translocations produce in-frame fusions of KMT2A to other chromatin regulatory proteins. Here we develop a strategy to map the genome-wide occupancy of oncogenic KMT2A fusion proteins in primary patient samples regardless of fusion partner. By modifying the versatile CUT&Tag method for full automation we identify common and tumor-specific patterns of aberrant chromatin regulation induced by different KMT2A fusion proteins. Integration of automated and single-cell CUT&Tag uncovers epigenomic heterogeneity within patient samples and predicts sensitivity to therapeutic agents.

## Introduction

Ten percent of acute leukemias harbor chromosomal translocations involving the *Lysine Methyl-transferase 2A* (*KMT2A*) gene (also referred to as *Mixed Lineage Leukemia-1*)^1^. In its normal role, KMT2A catalyzes methylation of the lysine 4 residue of the histone H3 nucleosome tail (H3K4) and is required for fetal and adult hematopoiesis^2^. The N-terminal portion of KMT2A contains a low complexity domain that mediates protein-protein interactions, an AT-hook/CXXC domain that binds DNA, and multiple chromatin-interacting domains (PHD domains and a bromo domain), whereas the C-terminal portion contains a trans-activation domain that interacts with histone acetyl-transferases and a SET domain that catalyzes histone H3K4 methylation^3,4^. The KMT2A pre-protein is cleaved to form a 320-kDa N-terminal fragment (KMT2A-N) and a 180-kDa C-terminal fragment (KMT2A-C) that form a stable dimer^5,6^.

*KMT2A* contributes to leukemogenesis through oncogenic chromosomal rearrangements involving the DNA-binding domain in the N-terminal portion of KMT2A with a diverse array of other chromatin regulatory proteins^7,8^. Although more than 80 translocation partners have been identified in *KMT2A*-rearranged (*KMT2A*r) leukemias, fusions involving AF9, ENL, ELL, AF4 or AF10 transcriptional elongation factors account for the majority of cases^1,8^. These fusion partners regulate RNA Polymerase II (RNAPII) elongation (ELL and AF4) or recruit the Dot1L-H3K79 histone methyltransferase (AF10), or both (AF9 and ENL)^9-12^. Additionally, ENL and AF9 interact with the CBX8 chromobox protein to neutralize the PRC1 gene silencing complex^13-16^.

Previous work has suggested that KMT2A fusion proteins bind different genomic loci depending on the fusion partner to drive different leukemia subtypes^17,18^. For example, AF4 fusions are more common in acute lymphoid leukemia (ALL), and AF9 fusions are associated with acute myeloid leukemia (AML)^1^. In addition, *KMT2A* rearrangements are also prevalent in mixed lineage leukemia (MPAL), and numerous examples of *KMT2A*r leukemias that interconvert between lineage types have been documented^17,19-21^. However, because methods for efficiently and reliably profiling KMT2A fusion binding sites in patients samples are lacking, the relationship between KMT2A fusions, chromatin structure, and lineage plasticity has been challenging to fully characterize. Here, we establish a chromatin profiling platform that efficiently profiles oncogenic fusion proteins, transcription-associated complexes, and histone modifications in cell lines and patient samples. By integrating these results with related single-cell methods we characterize the regulatory dynamics of *KMT2A*r leukemias. We identify groups of fusion oncoprotein target genes that show divergent patterns of active and repressive chromatin within the same sample. These patterns suggest that KMT2A-fusion proteins activate distinct oncogenic networks within different cells of the same tumor, and may explain lineage plasticity associated with *KMT2A*r leukemia. In addition, we find that distinct fusion partners display differential affinity for various transcriptional cofactors that predicts cancer sensitivity to therapeutic compounds.

## Results

### A strategy for mapping the binding sites of diverse KMT2A fusion proteins

Characterizing the chromatin localization of oncogenic fusion proteins has often been limited by the inability of ChIP-seq to be used with small amounts of patient samples. To efficiently compare the binding sites for wildtype KMT2A and the fusion proteins, we applied AutoCUT&RUN^22^ across a panel of four *KMT2A*r leukemia cell lines and eight primary *KMT2A*r patient samples sorted for CD45-positive blasts. This collection spans the spectrum of *KMT2A*r leukemia subtypes with diverse *KMT2A* translocations that create oncogenic fusion proteins with the transcriptional elongation factors AF4 (SEM, RS4;11, 1^0^ ALL-1 and 1^0^ MPAL-2), AF9 (1^0^ AML-3, 1^0^ MPAL-1), ENL (KOPN-8, 1^0^ AML-2), AF6 (ML-2), AF10 (1^0^ AML-4, 1^0^ AML-5), or a relatively rare fusion to the cytoplasmic GTPase Sept6 (1^0^ AML-1) (**Supplementary Table 1**). With the exception of ML-2, an AML-derived cell line, these samples also contain a wildtype copy of the *KMT2A* locus. For comparison, we also profiled KMT2A localization in untransformed human CD34+ hematopoeitic stem/progenitor cells (HSPCs), in H1 human embryonic stem cells, and in the K562 leukemia cell line, each of which lack *KMT2A* translocations. Antibodies to the C-terminal portion recognize only wildtype KMT2A-C, while antibodies to the N-terminal portion recognize both wildtype KMT2A-N and the fusion protein (**Fig. 1a**). Therefore, binding sites unique to the oncogenic fusion protein can be identified by comparing chromatin profiling of C-terminal and N-terminal KMT2A antibodies. We used AutoCUT&RUN to profile replicate samples of cell lines with two different antibodies to the N-terminus and two to the C-terminus of KMT2A, and correlation analysis of sequencing results showed high reproducibility (**Supplementary Fig. 1a**).

**Figure 1.**
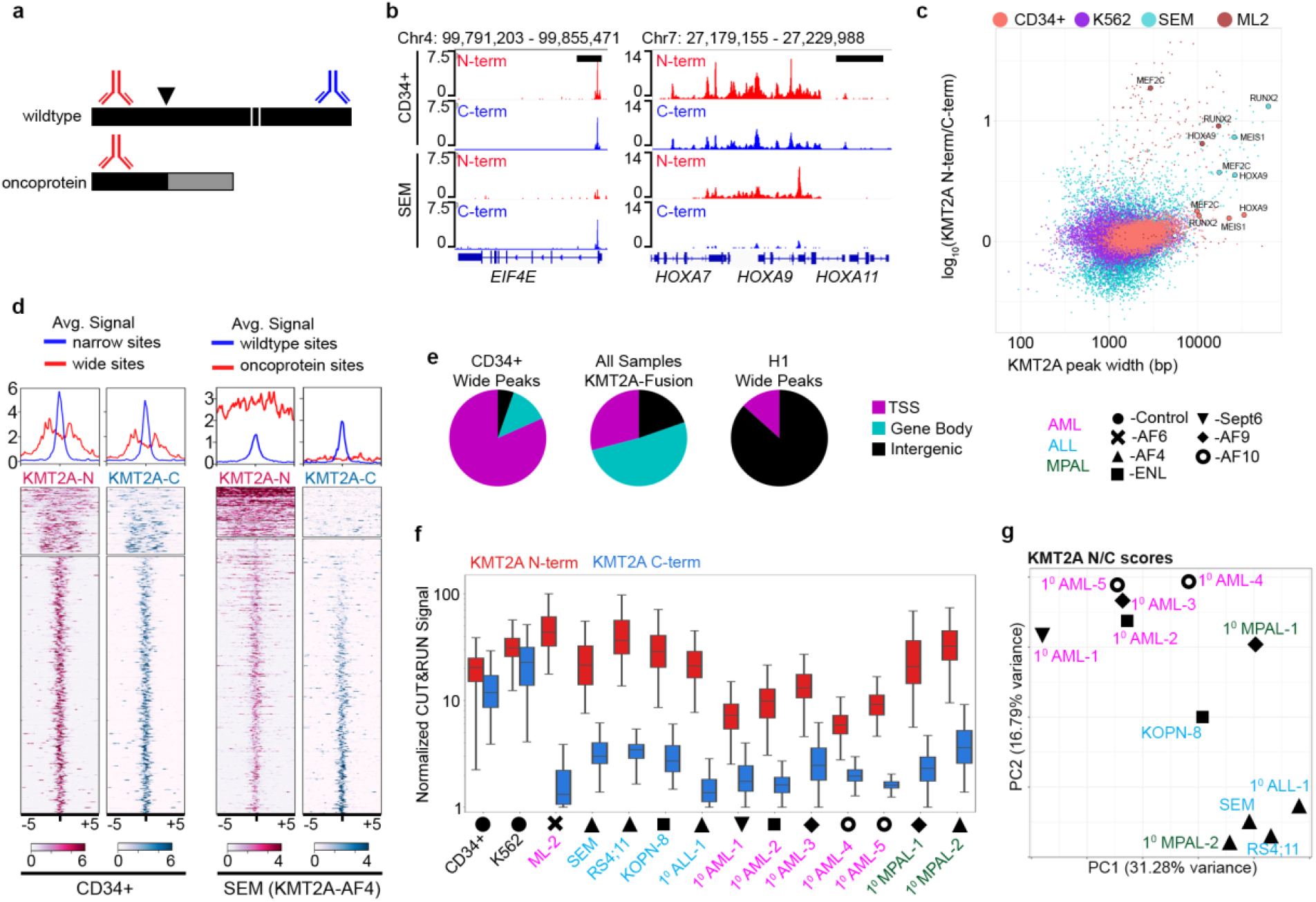
AutoCUT&RUN profiling of KMT2A-fusion protein binding. **a**, A general strategy for mapping KMT2A-fusion proteins. The wildtype KMT2A (black) is cleaved (white lines) into KMT2A-N and KMT2A-C proteins. Common oncogenic lesions (black arrowhead) produce in-frame translation of oncogenic KMT2A with numerous fusion partners (grey). C-terminal KMT2A antibodies (blue) recognize wildtype KMT2A-C. N-terminal KMT2A antibodies (red) recognize wildtype KMT2A-N and the oncogene KMT2A-fusion proteins. **b**, Example of a wildtype KMT2A binding site *(EIF4E)* and an oncoprotein binding site *(HOXA locus)*. Black scale bars = 10kb. **c**, KMT2A scatter plot comparing the peak width and relative enrichment of the KMT2A N-versus C-terminus in control (CD34+ and K562) samples and *KMT2A*r samples (SEM & ML-2). **d**, Heatmap comparison of KMT2A signal over broad CD34+ or KMT2A fusion binding sites. **e**, Pie chart of KMT2A-N signal at TSSs, gene bodies and intergenic regions shows KMT2A-fusion oncoproteins enrichment in gene bodies. **f**, Box plot of KMT2A-N and -C signal shows C-terminal KMT2A antibody depletion at KMT2A-fusion binding sites. **g**, PCA of fusion oncoprotein binding sites in *KMT2A*r samples. The first two components are shown. Statistics are included in Supplementary Table 3.

As expected, in H1, K562 and CD34+ HSPCs KMT2A-N and KMT2A-C show nearly identical patterns of enrichment across the genome (**Fig. 1b, Supplementary Fig. 1b**). Strikingly, in H1 cells KMT2A binding is generally focused in narrow peaks directly over transcriptional start sites (TSSs), whereas in K562 cells and CD34+ progenitors additional regions show wide peaks of both KMT2A-N and KMT2A-C extending from TSSs out across gene bodies. Many of the genes with wide KMT2A distribution in CD34+ progenitors are master regulators of hematopoietic cell fate, and have previously been defined as KMT2A fusion oncoprotein targets in leukemias^18,23^. To systematically define fusion protein binding sites across our collection of samples, we used Gaussian mixture modeling to partition KMT2A peaks into two different distributions based on both the width of KMT2A peaks and the enrichment-normalized ratio of KMT2A-N to KMT2A-C signal (KMT2A N/C score) (**Fig. 1c, Supplementary Fig. 2**). In the CD34+ HSPCs and K562 cells numerous sites are called as wide peaks and display log-transformed KMT2A N/C scores close to zero, indicating similar enrichment of both KMT2A-N and KMT2A-C proteins (**Fig. 1c Supplementary Fig. 2a)**. In contrast, wide peaks display high N/C scores in each of the *KMT2A*r leukemia samples, indicating enriched binding of KMT2A-N that we attribute to the fusion oncoprotein (**Fig. 1c, Supplementary Fig. 2b**). For example, in the SEM cell line 8,168 narrow peaks are identified with enrichment of both KMT2A-N and KMT2A-C, whereas 91 peaks are wide and enriched for KMT2A-N, which we interpret as fusion oncoprotein binding sites (**Fig. 1d, Supplementary Fig. 2b)**. Many of these sites have been previously identified^24,25^. Applying this strategy to the ML-2 cell line, which has a deletion of the wildtype *KMT2A* allele and only carries a KMT2A fusion oncoprotein, results in only 210 KMT2A-bound sites (**Fig. 1c, Supplementary Fig. 2c**). The majority of these sites are wide with a high KMT2A N/C score. The localization of the KMT2A fusion oncoprotein in ML-2 cells demonstrates that chromatin binding of fusion oncoproteins is not dependent on wildtype KMT2A protein, and validates our mapping approach.

Next, we examined how the distribution of KMT2A changes between the wildtype and oncofusion proteins. In CD34+ HSPCs 81% of wide peaks overlap a gene TSS, whereas in *KMT2A*r samples only 30% of fusion oncoprotein peaks overlap a TSS and 50% overlap a gene body (**Fig. 1e**). In comparison, in the control H1 human embryonic stem cell line only 15 of 17,000 KMT2A peaks were called as “wide” and 13 of those peaks fall in intergenic regions with KMT2A N/C scores close to zero (**Fig. 1e, Supplementary Figure 2a, Supplementary Figure 3a**). By comparing the enrichment of the KMT2A N-terminus and C-terminus across the fusion oncoprotein binding sites in all *KMT2A*r samples, we find that in all cases the N-terminus is more enriched than the C-terminus, and that this difference becomes exacerbated at the fusion oncoprotein binding sites due to a significant depletion of the C-terminal signal (**Fig. 1f**).

We then compared oncoprotein target sites between different leukemias with KMT2A-fusions (**Supplementary Figure 3c-m)**. We found that 81/440 (∼18%) of all fusion oncoprotein target genes are shared between five or more of the *KMT2A*r leukemia samples we profiled, representing 12% of the total sequence space occupied by the fusion proteins (**Supplementary Figure 3n,o**). As expected, the group of genes we identified as the most frequent KMT2A-fusion targets across our collection of samples is highly enriched for master regulators of hematopoiesis as well as genes that are required for *KMT2A*r leukemia^26-30^ (**Supplementary Table 2**). Principal Component Analysis (PCA) of KMT2A N/C scores across all oncoprotein binding sites indicates that both the specific fusion partner as well as the myeloid versus lymphoid lineage bias of the tumor may influence tumor-specific localization of the oncofusion protein (**Fig. 1g**). For example, all KMT2A-AF4 samples cluster together in the PCA plot and group with an ALL patient sample and one MPAL patient sample. In contrast the ALL cell line KOPN-8 which carries a KMT2A-ENL fusion protein partitions away from KMT2A-AF4-bearing leukemias. Primary AML samples bearing KMT2A-AF9, -AF10 and -ENL fusions form a second cluster, apart from the KMT2A-Sept6-containing primary AML and the primary KMT2A-AF9-bearing MPAL sample. Thus, tumors bearing KMT2A-AF4 fusions share a distinct binding profile, but other oncofusion proteins show both common and diverse localization patterns.

### Chromatin landscapes of *KMT2A*r leukemia samples

To economically characterize the global chromatin landscape of tumors at a scale that could be generally applied to patient samples we developed AutoCUT&Tag, a modification of our previous AutoCUT&RUN robotic platform^22^. CUT&Tag takes advantage of the high efficiency and low background of antibody-tethered Tn5 tagmentation-based chromatin profiling relative to previous methods, such as ChIP-seq and CUT&RUN^31^. The standard CUT&Tag protocol requires DNA extraction before library enrichment by PCR. However, we recently developed conditions for DNA release and PCR enrichment without extraction (CUT&Tag-direct)^32^. In this improved protocol a low concentration of SDS is used to displace bound Tn5 from tagmented DNA, and subsequent addition of the non-ionic detergent Triton-X100 quenches the SDS to allow for efficient PCR. This streamlined protocol makes CUT&Tag compatible with robotic handling of samples in a 96-well plate format and generates profiles with data quality comparable to those produced by benchtop CUT&Tag (**Supplementary Fig. 4**). To define the chromatin features around KMT2A fusion protein binding sites, we used AutoCUT&Tag to profile the active chromatin modifications Histone-3 Lysine-4 monomethylation (H3K4me1), Histone-3 Lysine-4 trimethylation (H3K4me3), Histone-3 Lysine-36 trimethylation (H3K36me3), Histone-3 Lysine-27 acetylation (H3K27ac), Histone-4 Lysine-16 acetylation (H4K16ac), and initiating RNA-Polymerase 2 marked by Serine-5 phosphorylation of the C-terminal domain (RNAP2S5p). In addition we profiled the silencing histone modifications Histone-3 Lysine-27 trimethylation (H3K27me3) and Histone-3 Lysine-9 trimethylation (H3K9me3). Together, these eight modifications distinguish active promoters, enhancers, transcribed regions, developmentally silenced, and constitutively silenced chromatin^33^, and provide a straightforward picture of the regulatory status of a genome (**Fig. 2a, Supplementary Fig. 4c**). Replicate profiles for each mark in control CD34+ samples and *KMT2A*r leukemia samples were very similar and were merged for further analysis (**Supplementary Fig. 5**). We first compared the chromatin features associated with sites bound by wildtype KMT2A or the KMT2A oncofusion protein across all samples. Consistent with the localization of the KMT2A-fusion proteins to actively transcribed genes, we found the active promoter marks H3K4me3, RNAP2S5p, and H3K27ac are all present at oncofusion protein binding sites (**Supplementary Figure 6a-c**). H3K4me3 is also enriched at some promoters in the ML-2 cell line (*e*.*g*., LPO and LYZ in **Figure 2b**), which lacks the KMT2A methyltransferase domain, indicating that another H3K4me3 methyltransferase is responsible. When compared to the sample-matched wildtype KMT2A-bound sites, H3K27ac is enriched at the oncofusion protein samples [SEM, KOPN-8, 1^0^AML-1 (a KMT2A-SEPT6 fusion), 1^0^AML-2, 1^0^AML-3, and 1^0^AML-5] (**Supplementary Figure 6b**). The H3K4me3 mark is significantly enriched at the oncofusion protein binding sites of five of the samples (SEM, RS4;11, 1^0^AML-1, 1^0^AML-2 and 1^0^MAPL-2), and significantly depleted in five of the other samples (1^0^ALL-1, 1^0^AML-3, 1^0^AML-4, 1^0^AML-5 and 1^0^MAPL-1) (**Supplementary Fig. 6c)**. Oncofusion protein binding sites lack H3K27me3 or H3K9me3 (**Supplementary Figure 6d**,**e**), but are enriched in H3K4me1 and H3K36me3, both of which mark transcribed gene bodies (**Supplementary Figure 6f**,**g**). Enrichment of these marks is expected for mis-targeting of KMT2A fusions into gene bodies^34^.

**Figure 2.**
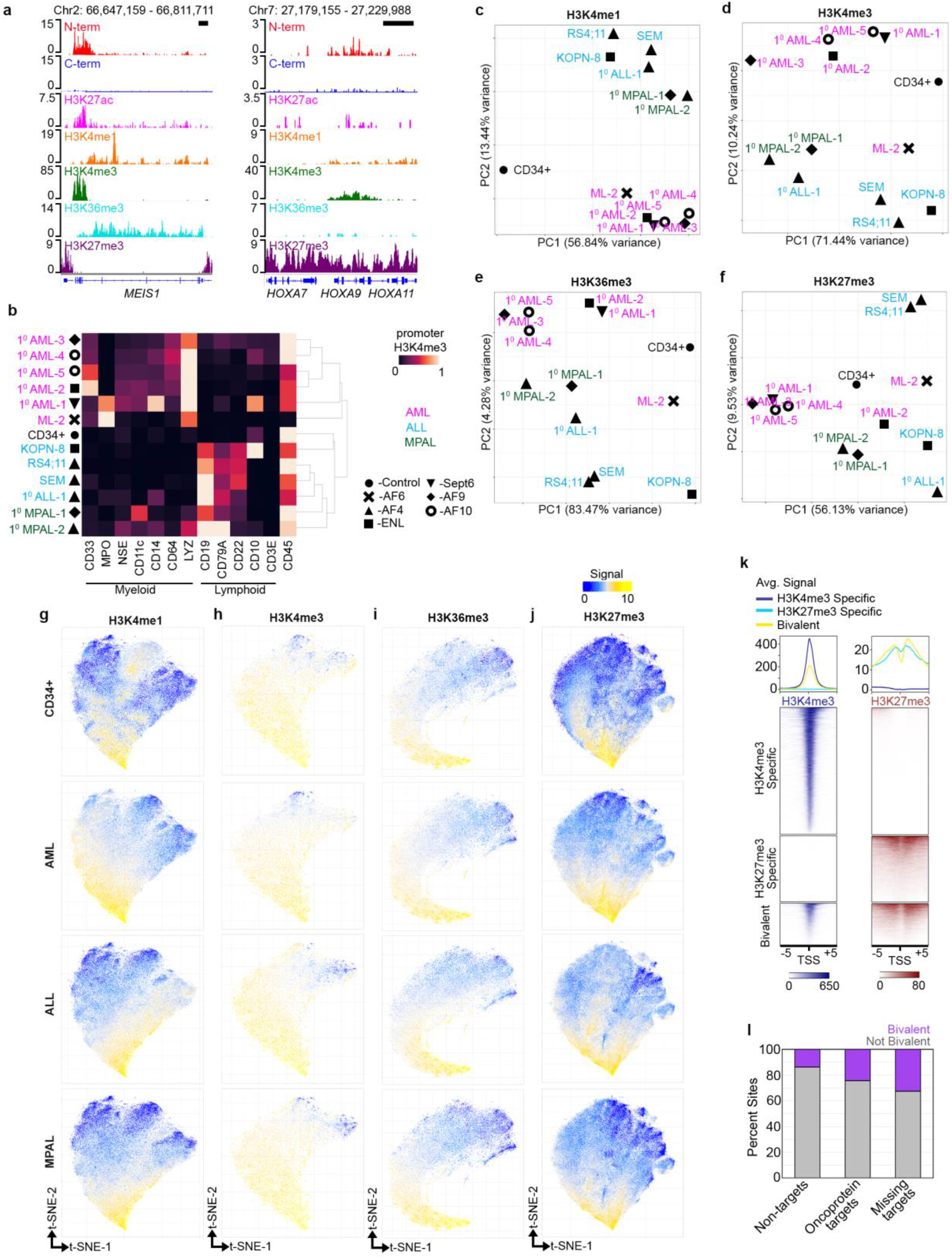
Clustering regulatory features distinguishes common and restricted elements in leukemia samples. **a**, the *MEIS1* locus is a direct target of KMT2A-AF9 in the 1^0^ MPAL-1 sample and is decorated by both active and repressive chromatin marks. The *HOXA* cluster is relatively repressed within this tumor. **c**, H3K4me3 signal at the promoters of diagnostic immunophenotypic markers accurately classifies AML, ALL and MPAL leukemias. **c**, PCA clustering analysis separates the H3K4me1-marked regions according to the lineage specificity. **d**, Same as (c) for H3K4me3. **e**, Same as (c) for H3K36me3. **f**, Same as (c) for H3K27me3. Grouping samples by PCA of their H3K27me3 repressive chromatin seperates tumors of the same lineage. **g**, Two-dimensional t-SNE projections separate lineage-specific H3K4me1 marked regions. Each colored pixel corresponds to a single H3K4me1 peak, colored by the maximum intensity within the indicated sample type. **h**, t-SNE projection of H3K4me3 identifies lineage specfic promoters. **i**, t-SNE plot of H3K36me3 marked regions in the indicated *KMT2A*r leukemia sugtypes. **j**, Same as (e) for H3K27me3. AML and ALL samples display H3K27me3 spreading, whereas H3K27me3 is more confined in the CD34+ control and the MPAL samples. **k**, Heatmap showing the regions called as bivalent in the 1^0^ MPAL-1 sample. **l**, Comparison of bivalency vs univalency at genes either not bound by the KMT2A fusion (non-target), bound in all samples (Oncoprotein target), or bound in all but one sample (Missing targets) shows that a bivalent chromatin signature is enriched at oncoprotein target genes.

Histone modification profiling holds the potential to reveal similarities and distinctions between leukemias by reporting their transcriptional status. For example, H3K4me3 reports gene promoter activity, and is enriched at marker genes that match the immunophenotypic characterization of each leukemia (**Fig. 2b**). To compare how the global distribution of these marks varies between *KMT2A*r leukemia samples, we first identified regions enriched for each modification in our collection of *KMT2A*r leukemia samples as well as CD34+ HSPCs using the SEACR peak-calling method^35^, and performed PCA to cluster samples according to their modification-specific similarities. Overall, active chromatin features marked by H3K4me1, H3K4me3, H3K36me3, H4K16ac, or RNAP2S5p cluster samples according to their ALL, AML, and MPAL lineage designation (**Fig. 2c-e, Supplementary Fig. 7a**,**b**), suggesting similar repertoires of active genes are used in each leukemia subtype. In contrast, PCA based on H3K27ac or H3K27me3 CUT&Tag profiles partitions samples into groups largely unrelated to their leukemia subtype (**Fig. 2f, Supplementary Fig. 7c**), and only the 1^0^AML-1 sample is distinguished by H3K9me3 (**Fig. 2f, Supplementary Fig. 7d**). H3K27me3 is an epigenetically inherited histone modification that is linked to developmental progression as cells determine their identities. Thus, these distinct H3K27me3 leukemia landscapes may relate to hematopoietic transitions that are defective in each tumor.

We next examined the lineage-specific variation in gene and regulatory element usage as indicated by the global chromatin landscape of each of the marks we profiled by performing t-distributed stochastic neighbor embedding (t-SNE) of these elements, followed by density peak clustering^36^. This analysis revealed that H3K4me1-marked regions are highly variable between lineage subtypes, with a substantial fraction of elements marked specifically in the AML samples falling to one side of the t-SNE plot, ALL specific elements partitioned to the other side of the plot, and CD34+ HSPC elements grouped in the middle (**Fig. 2g**). A fraction of both the AML and ALL specific elements are also marked by H3K4me1 in CD34+ cells and the primary MPAL samples we profiled (**Fig. 2g**). This regulatory overlap implies that MPAL leukemias share features with both ALL and AML, and that *KMT2A*r leukemia samples maintain H3K4me1 at regulatory elements employed during normal hematopoiesis.

In comparison to H3K4me1, a much larger fraction of H3K4me3 and H3K36me3 peaks are common across leukemia subtypes, indicating that they largely share gene expression repertoires (**Figs. 2h**,**i)**. Grouping H3K4me3-marked promoter regions by t-SNE also partitioned AML- and ALL-specific elements to opposing sides of the t-SNE graph, and identified groups of elements that are shared with the MPAL samples and CD34+ stem and progenitors (**Fig. 2h**), however as compared to H3K4me1, a smaller proportion of H3K4me3-marked features show any lineage specificity. This is consistent with previous reports that regulatory elements marked by H3K4me1 generally show more cell-type specificity than promoter elements marked by H3K4me3^37,38^.

Similar to the t-SNE analysis of H3K36me3 marked regions, t-SNE analysis of H3K27ac-, H4K16ac-, or H3K9me3-marked regions did not partition the genome by lineage identities (**Supplementary Fig. 7e-g**). In contrast, both the RNAP2S5p and H3K27me3 peaks showed diversity similar to H3K4me1 (**Fig. 2j, Supplementary Fig. 7h**). Analysis by t-SNE with H3K27me3 did not partition elements according to their lineage subtypes (**Fig. 2j**). Rather, AML and ALL samples have a greater proportion of the genome that is marked by H3K27me3 than in CD34+ cells, suggesting that they are more differentiated (**Fig. 2j**). Consistent with this interpretation, MPAL samples have fewer regions marked by H3K27me3, and are considered to have a higher degree of lineage plasticity (**Fig. 2j**). We conclude that high-throughput CUT&Tag profiling provides a powerful tool to characterize *KMT2A*r leukemias, and that profiling the developmentally repressed genome reveals tumor-specific differences that are not apparent by profiling the active genome.

### Bivalent chromatin signatures at KMT2A oncofusion protein target sites

In addition to marking promoters that are engaged in active transcription, H3K4me3 is present at a limited subset of transcriptional repressed, “bivalent” promoters that are also marked by H3K27me3^39,40^. In our collection of leukemia samples we observed both H3K4me3 and H3K27me3 at some promoters that are called as KMT2A-fusion protein targets (**Fig. 2a, left side**). Additionally, we observed genes that are bound by the oncofusion protein in the majority of *KMT2A*r leukemia samples, but are not called as targets in specific samples; we termed this group “missing targets” (**Fig. 2a, right side**). To systematically define the bivalent promoters within our collection of samples, we quantified the abundance of H3K4me3 and H3K27me3 within 2-kb windows centered on gene TSSs of marked and unmarked promoters for each modification. By intersecting these groups, we identified ∼2,000-5,000 bivalent promoters in each of the *KMT2A*r leukemia samples (**Fig. 2k, Supplementary Fig. 8**). Interestingly, we found that ∼33% (129/396) of the missing target promoters are called as bivalent, whereas ∼24% (267/1097) of KMT2A-fusion target promoters are bivalent, and only 14% of wildtype KMT2A targets are bivalent (**Fig. 2l**). Thus, oncofusion protein target promoters are enriched for a bivalent chromatin signature, suggesting that expression of these genes may fluctuate between cells within a sample.

### Single cell CUT&Tag reveals heterogeneity of bivalent KMT2A-fusion target genes

To test if bivalent histone modifications at KMT2A-fusion target promoters are due to heterogeneity between cells, we performed single-cell CUT&Tag on 8 *KMT2A*r leukemia samples. Antibody binding and pA-Tn5 tethering were performed on bulk samples and then individual cells were arrayed in microwells on the ICELL8 platform for barcoded PCR library enrichment^31^. We optimized the median number of unique reads per cell while maintaining a high fraction of reads in peaks on the ICELL8 by varying the amount of SDS detergent to release Tn5 after tagmentation and of Triton X-100 to quench SDS before PCR (**Supplementary Fig. 9a**,**b**). Using this approach, we profiled 1,137-3,611 cells for the H3K4me3, H3K27me3, and H3K36me3 histone modifications. After excluding cells with <300 fragments, single-cell CUT&Tag for H3K4me3, H3K27me3, and H3K36me3 yielded medians of 4,972, and 13,025, and 3,962 unique reads per cell, respectively (**Supplementary Fig. 9c**). As a second quality control step, we called peaks on the aggregate data of all cells profiled for each mark and removed cells that had a fraction of reads in peaks (FRiP) below the normal distribution (**Supplementary Fig. 9d**,**e**). Profiles for each single cell were then split into 5-kb bins tiled across the genome and cells were projected in UMAP space based on that binning profiling, but not on H3K36me3 (**Fig. 3a-c**). This implies that the leukemia samples differ in both sets of active promoters and in silenced regions.

**Figure 3.**
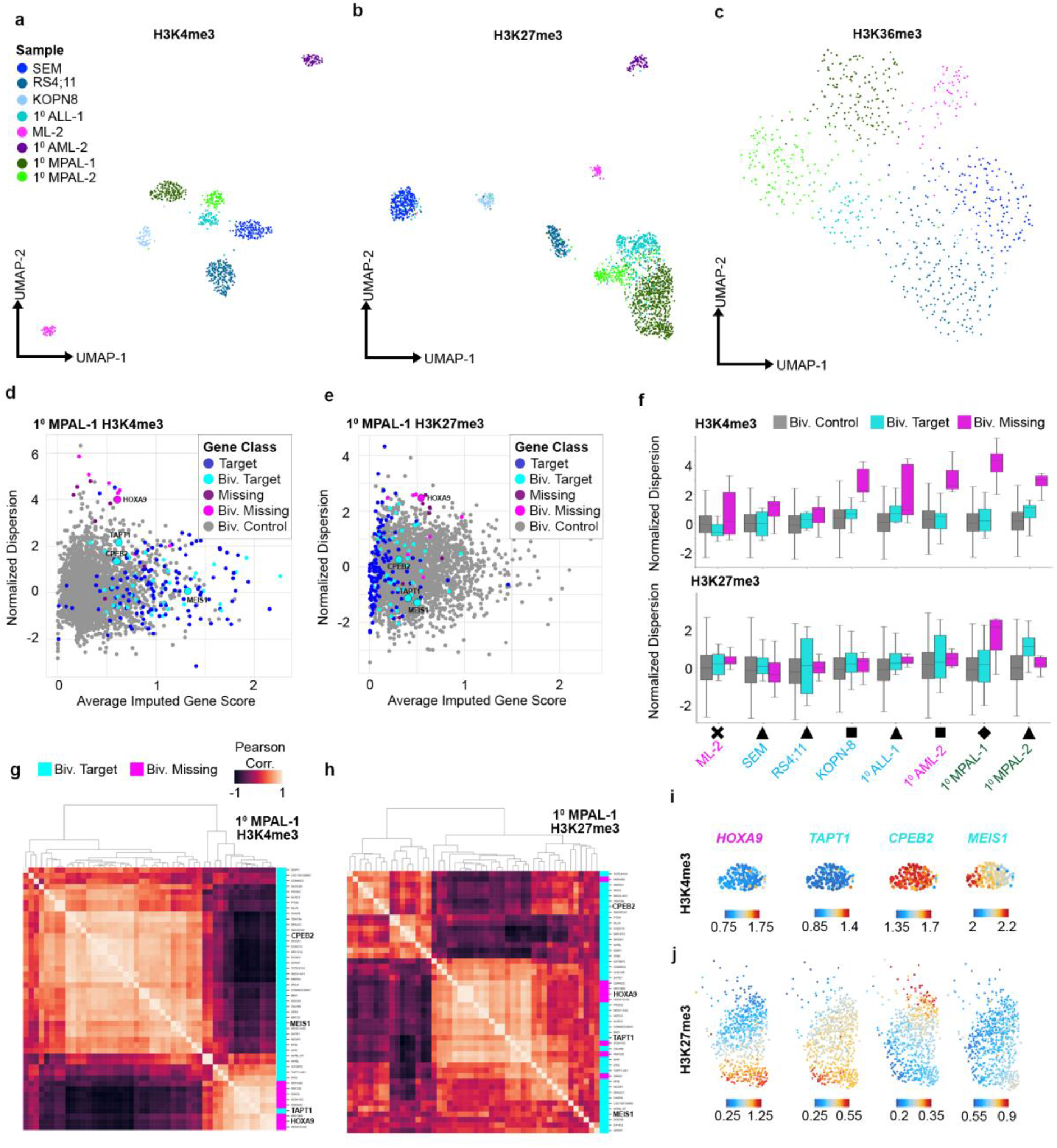
Single Cell profiling of H3K4me3 and H3K27me3 reveals chromatin heterogeneity at KMT2A-fusion target loci. **a**, UMAP projection of the H3K4me3 profiles in single leukemia cells resolves sample specific clusters. The H3K4me3 count matrix for each cells split reads into 5 kb bins tiled across the genome. **b**, Same as (a) for H3K27me3. A fraction of cells in the 1^0^ ALL-1, 1^0^ MPAL-1 and 1^0^ MPAL-2 cells intermingle in H3K27me3 UMAP space. **c**, Same as (a) for H3K36me3. Leukemia cells do not form tight sample specific groups according to their H3K36me3 profiles. **d**, Scatterplot comparing the 1^0^ MPAL-1 average imputed H3K4me3 scores and normalized dispersion of genes grouped according to the KMT2A-fusion binding status (Target, Missing Target, and unbound Control) and promoter bivalency status (Biv.) in bulk profiling assays. Select genes are highlighted. **e**, Same as (d) for H3K27me3. **f**, KMT2A-fusion targets show elevated H3K4me3 and H3K27me3 dispersion across select leukemia samples. **g**. Organizing genes according to the co-variance of H3K4me3 imputed gene scores across 1^0^ MPAL-1 cells resolves groups that vary in concert with one another but are anti-correlated with genes in the other group. Selected genes are highlighted for comparison. **h**, Same as (g) for H3K27me3. **i**, Imputed H3K4me3 gene scores of the highlighted genes from (g) and (h) displayed on the UMAP of 1^0^ MPAL-1 cells from (a). **j**, Same as (i) for H3K27me3. *HOXA9* and *TAPT1* have high H3K4me3 scores in a limited subset of the tumor, but high H3K27me3 scores in the majority of the tumor, whereas *CPEB2* and *MEIS1* show the opposite pattern and are enriched for H3K4me3 in the majority fo the tumor, and rarely show high H3K27me3 scores. Statistics are included in Supplementary Table 3.

To examine intra-tumor heterogeneity in the H3K4me3 and H3K27me3 signals we first used the archR single-cell software package^41,42^ to calculate imputed gene scores for all genes according to the UMAP projection of all cells. We then determined the normalized dispersion of the imputed scores within cells of the same sample (**Fig. 3d, Supplementary Fig. 10**). Strikingly, bivalent missing targets show higher H3K4me3 dispersion in the SEM, KOPN-8, 1^0^ AML-2, 1^0^ MPAL-1 and 1^0^ MPAL-2 samples than in tumor-matched controls (**Fig. 3f**). This implies that the expression of these genes varies between cells.

Next, we examined variation in the repressive H3K27me3 mark at bivalent oncoprotein target genes. In 1^0^ MPAL-1 the normalized dispersion of H3K27me3 is higher in bivalent missing target genes, and in 1^0^ MPAL-2 the normalized H3K27me3 dispersion is higher in bivalent target genes (**Fig. 3f**). Some bivalent genes vary between cells for both the H3K4me3 and the H3K27me3 modifications. For example, the HOXA9 gene is a “missing target” in 1^0^ MPAL-1 cells (**Fig. 2a**), but shows high dispersion in both H3K4me3 and H3K27me3 signals (**Fig. 3d-e**). Thus, bivalency of chromatin marks is associated with heterogeneity between cells within a sample.

Grouping bivalent target genes by their similarity in imputed gene scores separates two groups by either H3K4me3 or H3K27me3 profiling (**Fig. 3g-h, Supplementary Fig. 11**). For example, the missing target gene *HOXA9* is enriched for H3K4me3 scores in only a small number of 1^0^ MPAL-1 leukemia cells (**Fig. 3i**). As expected from their similar gene scores (Fig. 3g), the *TAPT1* gene has the highest H3K4me3 scores in the same cells as *HOXA9* (**Fig. 3i**). In contrast, genes that are anti-correlated with *HOXA9* such as *CPEB2* and *MEIS1* (Fig 3g) have the weakest H3K4me3 signal in cells where *HOXA9* is active (**Fig. 3i**). This suggests that there are two exclusive gene expression programs activated by the KMT2A fusion oncoprotein. Furthermore we find the imputed H3K27me3 scores also form inverse patterns of gene association from H3K4me3, where genes with little H3K4me3 are highly enriched for H3K27me3 in the majority of tumor cells, whereas genes with high H3K4me3, rarely show H3K27me3 (**Fig. 3j**). These groups of divergent KMT2A fusion oncoprotein targets may contribute to the phenotypic plasticity of *KMT2A*r leukemias.

### Chromatin profiling for transcriptional cofactors predicts drug sensitivity of leukemias

We reasoned that the distinct binding sites of KMT2A fusion proteins may be driven in part by the cofactors with which the fusion oncoproteins associate. Therefore, we used AutoCUT&Tag to map the distributions of ENL and Dot1L, two chromatin proteins that interact with KMT2A fusion proteins^34^. Regions bound by KMT2A fusion proteins are enriched for Dot1L and ENL in all samples as compared to sample-matched wildtype KMT2A-bound sites (**Fig. 4a-c**). Dot1L has been proposed to be a central component of oncogenic transformation by KMT2A fusion proteins in certain leukemias^12,43^, and we find the Dot1L histone methyltransferase is enriched more at the oncofusion protein targets in KOPN-8, 1^0^AML-3 and 1^0^MPAL-1 samples than in the other leukemias we profiled (**Fig. 4b**). Both 1^0^AML-3 and 1^0^MPAL-1 carry a KMT2A-AF9 fusion protein, whereas KOPN-8 carries a KMT2A-ENL fusion, suggesting that the AF9 fusion partner has a particularly high affinity for Dot1L, while other leukemias can be variable for Dot1L recruitment. This is illustrated by the MLL-SEPT6 fusion (1^0^AML-1), where only modest enrichment of Dot1L and ENL at fusion binding sites was observed.

**Figure 4.**
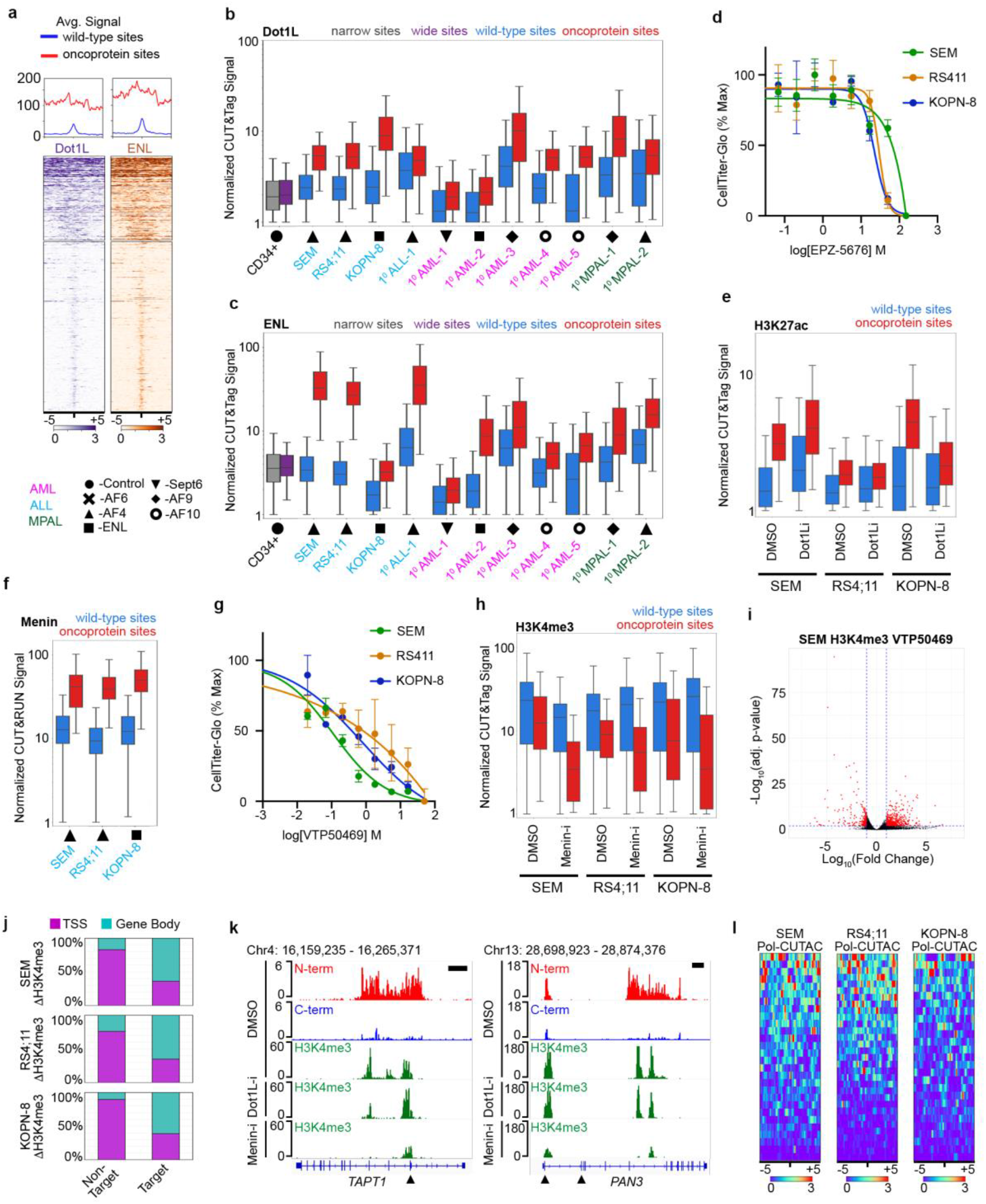
AutoCUT&Tag profiling reveals therapeutic sensitivities of *KMT2A*r leukemia samples. **a**, Heatmap comparing the co-ocupancy of the transcriptional cofactors Dot1L and ENL of the fusion oncoprotein sites (top) and sample-matched wildtype sites (bottom) in the 1^0^ MPAL-1 sample. **b**, Dot1L is significantly enriched at the fusion oncoprotein binding sites in all *KMT2A*r leukemia samples. The KMT2A-AF9 bearing 1^0^ AML-3 and 1^0^ MPAL-1 samples as well as the KMT2A-ENL cell line KOPN-8 show the highest degree of enrichment of Dot1L and fusion oncoprotein sites. **c**, ENL is significantly enriched at the fusion oncoprotein binding sites in all *KMT2A*r leukemia samples. **d**, Cell survival curves of SEM, RS4;11 and KOPN-8 cell lines in response to increasing concentrations of the Dot1L inhibitor EPZ-5676. **e**, In response to treatment with 30 µM of the Dot1L inhibitor for three days, H3K27ac is depleted at the fusion oncoprotein binding in KOPN-8 cells but not SEM or RS4;11.**f**, The transcriptional scaffold protein Menin is enriched at the fusion oncoprotein binding sites to a similar extent in SEM, RS4;11 and KOPN-8 cells. **g**, Same as (d), but treated with the inhibitor of KMT2A-Menin binding VTP50469 **h**, In response to treatment with 30 µM of the Menin inhibitor for three days, H3K4me3 is depleted at the fusion oncoprotein binding in samples. **i**, Volcano plot identifying SEM H3K4me3 peaks where the signal changes in response to Menin inhibition. **j**, Comparison of H3K4me3 signal at TSSs and gene bodies shows that loss of H3K4me3 methylation occurrs preferentially in the gene bodies of KMT2A-fuision target genes. **k**, Two examples of KMT2A-AF4 bound genes in SEM cells that *(TAPT1 and PAN3)*, that lose H3K4me3 signal in the gene body in response treatment with 30 nM of the Menin inhibitor for 3 days. Black arrowheads point to annotated TSSs, Black scale bars = 10kb. **l**, Heatmaps showing the Menin sensitive H3K4me3 peaks located in the fusion oncoprotein target gene bodies of the indicated cell types are accessible and enriched for inititating RNA-Pol2, as indicated by RNAP2S5p CUTAC (Pol-CUTAC). Heatmaps are centered over the H3K4me3 peaks. Statistics are included in Supplementary Table 3.

Several studies have suggested that KMT2A-AF9 fusion leukemias are particularly sensitive to pharmacological inhibition of the Dot1L methyltransferase activity^44,45^. We hypothesize that elevated Dot1L signal at dprotein target sites might indicate sensitivity Dot1L inhibitors. Indeed, we found that KOPN-8 cells are more sensitive to the Dot1L inhibitor EPZ-5676 than either SEM or RS4;11 cells (**Fig. 4d**). Previous reports have also shown that KOPN-8 cells are sensitive to the Dot1L inhibitor EPZ-0007477^45^. Using AutoCUT&Tag to profile H3K27ac after drug treatment, we found a significant depletion of this histone mark at oncofusion protein binding sites, whereas H3K27ac at KMT2A-AF4 bound sites in SEM and RS4;11 leukemias remained unchanged (**Fig. 4e**). Thus, this pharmocological agent specifically alters the chromatin of oncofusion protein targets in *KMT2A*r leukemia samples.

To extend this analysis we profiled the transcriptional scaffold protein Menin which interacts with the N-terminal portion of KMT2A and with oncofusion proteins by AutoCUT&RUN. SEM, RS4;11 and KOPN-8 cells have similar levels of Menin at KMT2A-fusion bound sites, but the SEM cell line was more sensitive to the Menin inhibitor VTP50469 (**Fig. 4g**). We then used autoCUT&Tag to profile H3K4me3 in Menin-inhibited cells (**Fig. 4h**). We called H3K4me3 peaks that showed a depleted signal after drug treatment, and these depleted sites are highly enriched in the gene bodies of oncofusion protein targets (**Fig. 4i-k**). Finally, we examined chromatin accessibility and the presence of initiating RNA polymerase II (RNAP2S5p in drug-treated cells using Pol-CUTAC^46,47^, and found many oncofusion- and Menin-bound sites are normally highly accessibile and bound by intiating RNA Polymerase II (**Fig. 4l**). This supports the idea that oncofusion protein-induced transcription in *KMT2A*r leukemias is highly sensitive to Menin inhibition.

## Discussion

Here we have applied high-throughput chromatin profiling to *KMT2A*r leukemias to delineate fusion protein-specific targets and to identify chromatin features that are characteristic of myeloid, lymphoid and mixed-lineage leukemias. To economically profile these features we took advantage of the high signal-to-noise and low sequencing depth requirements inherent to CUT&RUN and CUT&Tag for full automation on a standard liquid handling robot. As CUT&Tag requires only thousands of cells for informative histone modifications^46^, AutoCUT&Tag is suitable for profiling of samples for a wide range of studies, including developmental and disease studies and screening patient samples. The enhanced throughput and consistency of the AutoCUT&RUN and AutoCUT&Tag platforms for chromatin profiling makes these technologies suitable for profiling patient specimens.

By also performing AutoCUT&RUN on KMT2A fusions and components of the SuperElongation and DotCom complexes we have elucidated mechanistic details that likely contribute to the heterogeneity of these tumors. We found the most common KMT2A-fusion proteins, including KMT2A-AF4, KMT2A-AF9 and KMT2A-ENL all colocalize with the ENL protein in gene bodies, whereas a relatively rare KMT2A-Sept6 fusion protein does not colocalize with ENL and also tends to be more tightly associated with promoters. This suggests that the interaction of the C-terminal domain of AF4, ENL and ELL with transcriptional elongation complexes likely recruits the fusion protein from the promoter into the gene-body. Consistent with the possibility that these interactions play a pivotol role in oncogenic transformation, the wildtype ENL allele is required for tumor growth in numerous *KMT2A*r cell lines^48^. In contrast, the cytoskeletal GTPase SEPT6 is not known to interact with SEC or DotCom components and instead may contribute to oncogenesis by promoting multimerization of the oncofusion protein.

Using AutoCUT&Tag to profile histone modifications of leukemia samples we identified frequent KMT2A fusion oncoprotein sites showing bivalent chromatin features. At some sites bivalent chromatin features correlated with heterogeneity between cells, implying that these populations are mixtures of cells with different gene expression programs. This has implications for how resistance to therapies may develop, if only a subset of cells are susceptible to specific anti-cancer agents.

Heterogeneity in leukemias may arise if an early cancerous cell divides and differentiates into two related cell types. Alternatively, certain leukemias may sporadically switch between cell types^24,25^. Our single cell profiling reveals that some leukemias display at least two gene expression programs that differ between individual cells. Kinetic analysis of chromatin dynamics within cell populations will be needed to determine whether bivalency reflects differentiation or sporadic switching, with implications for therapeutic strategies to limit relapse. Multiple compounds targeting chromatin proteins show promise as therapeutics for certain leukemias^45,49^. Profiling the targets of these compounds distinguishes certain *KMT2A*r leukemias in which Dot1L is enriched at KMT2A fusion oncoprotein target sites, thus providing a strategy for selecting patients suitable for treatment with Dot1L-targeting compounds. We also identified samples where KMT2A fusion oncoprotein target genes are broadly enriched for H3K4me3 in gene bodies and also bound by initiating RNAPII. These leukemias are particularly sensitive to treatment with the Menin inhibitor VTP50469, and again demonstrate the utiity of chromatin profiling for selecting therapeutic treatments. Incorporating AutoCUT&RUN and AutoCUT&Tag into longitudinal clinical trials could thus provide a route to assess the efficacy of epigenetic medicines. As these technologies are scalable and cost effective, the information obtained from chromatin profiling could be used for patient diagnosis.

### Data Accession

Gene Expression Omnibus GSE159608

## Author contributions

DHJ and SH optimized the CUT&Tag method for automation, and DHJ adapted these modifications for single cell CUT&Tag profiling. SM provided clinical samples and helpful discussion. DHJ, JFS, KA, and SH designed experiment. DHJ, EB and JFS performed experiments. DHJ, MPM, SJW and JFS performed data analysis. DHJ, MPM, KA, and SH wrote the manuscript. All authors read and approved the final manuscript.

## Acknowledgements

We thank the Fred Hutchinson Genomics Shared Resource Facility for technical support, particularly Phil Corrin and Jeff Delrow for help with AutoCUT&RUN profiling of KMT2A. We thank Terri Bryson and Trizia Llagas for help with cell culture, and Jorja Henikoff and Matt Fitzgibbon for preparing the sequencing data for analysis. In addition, we thank Jitendra Thakur and Scott Furlan for helpful discussions related to data analysis and presentation. We thank Charles Mullighan, Rhonda Ries, Jenny Lill and Marie Bleakley for generously sharing the *KMT2A*r samples and cell lines used in this study. This work was supported by NIH grants R01 HG010492 (S.H.), 4DN TCPA A093 (S.H.) and F32 GM129954 (M.M.), by the Howard Hughes Medical Institute (S.H.), by a pilot project grant from the Chan-Zuckerberg Initiative (S.H.), by a Damon Runyon-Sohn Foundation Fellowship (J.F.S.) and by an Alex’s Lemonade Stand Foundation Young Investigator Award (J.F.S.).

## Methods

### Cell Culture

Human K562 cells were purchased from ATCC (Manassas, VA, Catalog #CCL-243) and cultured according to supplier’s protocol. H1 hESCs were obtained from WiCell (Cat# WA01-lot# WB35186) and cultured in Matrigel™ (Corning) coated plates in mTeSR™1 Basal Media (STEMCELL Technologies cat# 85851) containing mTeSR™1 Supplement (STEMCELL Technologies cat# 85852). The *KMT2A*r cell lines ML-2, KOPN-8, RS4;11 and SEM were obtained from the Bleakley lab at the Fred Hutchinson Cancer Research Center.

### Drug Treatment

10,000 SEM, RS4;11, and KOPN-8 were plated in 90 µL of the appropriate media (see above) in a 96-well cell culture plate. A serial dilution of the Dot1L inhibitor EPZ-5676 and the Menin inhibitor VTP50469 were prepared in DMSO, and then diluted in primary media to control for the concentration of DMSO across all conditions. Ten µL of the diluted inhibitors was then added to the cell culture suspensions and mixed. Cells were then grown for 3 days at which point the viability was measured using a CellTiter-Glo assay (Promega Cat# G9241) read out on a standard luminometer. For chromatin profiling experiments SEM, RS4;11 and KOPN-8 cells were plated at the same density (10,000 cells/100 µL) in 20mL of media containing either 30 µM EPZ-5676, 30 µM VTP50469 or DMSO alone. After 3 days in cluture the cells were harvested and prepared for either AutoCUT&RUN or AutoCUT&Tag processing.

### Primary Patient Samples

Cryopreserved CD45 leukmia blasts for primary MPAL-1 (Sample ID: SJMPAL012424_D1, Alias TB-11-3295) and primary ALL-1 (Sample ID: SJALL048347_D1, Alias TB-13-0939) were obtained from St. Jude Children’s Research Hospital in accordance with institutional regulatory practices. Cryopreserved CD45 leukmia blasts for primary AML-1 (Sample ID: A40725), primary AML-2 (Sample ID: A67194) and primary MPAL-2 (Sample ID: A58548) were obtained from the Meshinchi lab at the Fred Hutchinson Cancer Research Center. Diagnosis of clinical samples as ALL, AML or MPAL was based on flow cytometry of samples stained with CD45-APC-H7 (BD Cat# 560178), cytoplasmic CD3-PE (BD Cat# 347347), CD34-PerCP Cy5.5 (BD Cat# 347203), CD18-APC (BD Cat# 340437), cytoplasmic MPO-FITC (Dako Cat# F071401-1), and CD33-PE-Cy7 (BD Cat# 333946). The KMT2A fusion present in each sample was determined by RNA-sequencing. The CD34+ hematopoietic stem and progenitor cells were obtained from the Fred Hutch Cooperative Centers of Excellence in Hematology Core in accordance with institutional regulatory practices.

### Antibodies

For profiling the wildtype and oncogenic KMT2A protein we used two monoclonal antibodies targeting the KMT2A N-terminus: Mouse anti-KMT2A (1:100, Millipore Cat #05-764) refered to as KMT2A-N1, and Rabbit anti-KMT2A (1:100, Cell Signaling Tech Cat #14689S) refered to as KMT2A-N2; as well as two monoclonal anitbodies targeting the KMT2A C-terminus: Mouse anti-KMT2A (1:100, Millipore Cat #05-765) refered to as KMT2A-C1, and Mouse anti-KMT2A (1:100, Santa Cruz Cat #sc-374392) refered to as KMT2A-C2. Since pA-MNase does not bind efficiently to many mouse antibodies, we used a rabbit anti-Mouse IgG (1:100, Abcam Cat# ab46540) as an adapter; this antibody was also used in the absence of a primary antibody as the IgG negative control. For profiling Menin via AutoCUT&RUN we used Rabbit anti-Menin (Bethyl Cat# A300-105A). For profling the SEC and Dotcom components via manual and and AutoCUT&Tag we used rabbit anti-ENL (Cell Signaling Tech Cat# 14893S), rabbit anti-ELL (Cell Signaling Tech Cat# 14468S) and rabbit anti-Dot1L (Cell Signaling Tech Cat# 90878S). For profiling histone marks via manual and AutoCUT&Tag, as well as single-cell CUT&Tag we used Rabbit anti-H3K4me1 (1:100 Thermo Cat# 710795), Rabbit anti-H3K4me3 (1:100 for bulk profiling or 1:10 for single-cell experiments, Active Motif Cat# 39159), Rabbit anti-H3K36me3 (1:100 for bulk profiling or 1:10 for single-cell experimemts, Epicypher Cat# 13-0031), Rabbit anti-H3K27me3 (1:100 for bulk profiling or 1:10 for single-cell experimetns, Cell Signaling Technologies Cat# 9733S), Rabbit anti-H3K9me3 (1:100, Abcam Cat# ab8898), Rabbit anti-H3K27ac (1:50 Millipore Cat# MABE647), Rabbit anti-H4K16ac (1:50, Abcam Cat# ab109463), and Rabbit anti-RNAPIISer5P (1:100, Cell Signaling Technologies Cat# 13523). To increase the local concentration of pA-Tn5, all CUT&Tag reactions also included the secondary antibody Guinea Pig anti-Rabbit IgG (1:100, antibodies-online Cat# ABIN101961).

### AutoCUT&RUN

Primary patient samples were thawed at room temperature, washed and bound to Concanavalin-A (ConA) paramagnetic beads (Bangs Laboratories Cat# BP531) for magnetic separation. Samples were then suspended in Antibody Binding Buffer and split for incubation with either the KMT2A N- or C-terminus specific antibodies or the IgG control antibody overnight. Sample processing was performed by the CUT&RUN core facility at the Fred Hutchinson Cancer Research Center according to the AutoCUT&RUN protocol available through the Protocols.io website (dx.doi.org/10.17504/protocols.io.ufeetje).

### CUT&Tag

Manual CUT&Tag reactions were performed according to the CUT&Tag-Direct protocol^32^. Briefly, nuclei were prepared by suspending cells in NE1 Buffer (20 mM HEPES-KOH pH 7.9, 10 mM KCl, 0.5mM Spermidine, 0.1% Triton X-100, 20% Glycerol) for 10 min on ice. Samples were then spun down and resuspended in Wash Buffer (20 mM HEPES pH 7.5, 150 mM NaCl, 0.5 mM Spermidine, Roche Complete Protease Inhibitor EDTA-Free) and lightly cross-linked by addition of 16% fomaldehyde to 0.1%. After 2 min, cross-linking was stopped by addition of 2.5 M glycine to a final concentration of 75 mM. Nuclei were washed and either cryopreserved in a Mr. Frosty Chamber for long term storage, or bound to ConA magnetic beads for further processing. ConA-bound nuclei were suspended in Antibody Binding Buffer (Wash Buffer containing 2 mM EDTA) and split into individual 0.5 mL tubes for antibody incubation at room temperature for 1 hr or 4°C overnight. Samples were then washed to remove unbound primary antibody, brought up in Wash buffer containing the secondary antibody, and incubated at 4°C for 1 hr. Samples were then washed and brought up in 300-Wash Buffer (Wash Bufer with 300 mM NaCl), containing pA-Tn5 (1:150 dilution), and incubated at 4°C for 1 hr. Samples were then washed in 300-Wash Buffer, and brought up in Tagmentation Buffer (300 Wash Buffer plus 10 mM MgCl2), and incubated at 37°C for 1 hr to allow the Tn5 tagmentation reaction to go to completion. Samples were then washed with TAPS wash buffer (10 mM TAPS with 0.2 mM EDTA), and brought up in 5 µL of Release Solution (10 mM TAPS with 0.1% SDS). Samples were then incubated in a thermocycler with heated lid at 58 degrees for 1 hr to release Tn5 and prepare tagmented chromatin for PCR. Neutralizing Solution (15 µL 0.67% Triton-X100) was added followed by 2 µL barcoded i5 primer (10 µM), 2 µL barcoded i7 primer (10 µM) and 25 µL of NEBNext PCR mix. Samples were then placed in a thermocycler and PCR amplification was performed using 12-14 rapid cycles. CUT&Tag libraries were then cleaned up with a single round of SPRIselect beads at a 1.3:1 v/v ratio of beads to sample, quantified on a Tapestation bioanalyzer instrument and pooled for sequencing.

### AutoCUT&Tag

A detailed protocol complete with program downloads has been made publicly available on protocols.io for implementing AutoCUT&Tag on a Beckman Coulter Biomek liquid handling robot (https://www.protocols.io/view/autocut-amp-tag-streamlined-genome-wide-profiling-bgztjx6n). To facilitate adaptation of the method to other standard liquid handling modules, the complete specifications for each step in the automated procedure are outlined in guidelines section. Briefly, nuclei were extracted, lightly cross-linked, bound to ConA beads and incubated with primary antibody as in manual CUT&Tag. Up to 96 samples were then arrayed in a 96 well PCR plate and positioned on a a stationary ALP on the Beckman Coulter Biomek FX Robot equipped with an ALPAQUA Magent Plate for standard magnetic separation, an ALPAQUA LE Magent Plate for low volume elution, and a thermal block for temperature controlled inbuation. Wash Buffer and 300-Wash Buffer were loaded in Deep Well Plates, Secondary Antibody Solution, pA-Tn5 solution, Tagmentation Buffer, TAPS Buffer and Release Buffer were all loaded into V-Bottom Plates and were positioned on Stationary ALPs in accordance with the preprogrammed AutoCUT&Tag method. The AutoCUT&Tag processing was conducted over the course of 4 hours. The sample plate containing ConA-bound tagmented nuclei in 10 µL 0.1% SDS was then removed, sealed and placed on a thermocycler with heated lid for a 1 hour incubation at 58°C. Using a reservoir and multichannel pipettor, 54 µL of 0.15% SDS neutralization solution was added to each well, followed by 4 µL of premixed i5/i7 barcoded primers, and 36 µL of premixed KAPA PCR Master Mix. The plate was then sealed and returned to a thermocycler for 14 rapid PCR cycles. Following PCR amplification, the sample plate was returned to the Biomek for one round of post-PCR cleanup on the Biomek deck set up in accordance a preprogrammed post-PCR cleanup method, including a second 96-well plate preloaded with SPRISelect Ampure beads, a Deep Well Plate loaded with 80% Ethanol for bead washes, and two V-Bottom Plates preloaded with 10 mM Tris-HCl pH 8.0 for tip washes and elution. Upon completion of the 1 hr cleanup the samples were then quantified using a Tapestation bioanalyzer instrument and pooled for sequencing.

### Single-cell CUT&Tag

Nuclei were extracted and lightly cross-linked using the same strategy as for manual CUT&Tag. The nuclei concentration was then quantified to allow for accurate dilution prior to dispensing into nanowells on the ICELL8. For each antibody 10 µL of ConA beads were washed in Binding Buffer (20 mM HEPES-KOH pH 7.9, 10 mM KCl, 1 mM CaCl_2_, 1 mM MnCl_2_) and bound to the sample for 10 min. Samples were the split into 0.5 mL Lobind tubes, one for each antibody, and resuspended in 25 µL of Antibody Buffer containing primary antibody at a 1:10 dilution. Samples were incubated at 4°C overnight, washed twice with 100 µL of Wash Buffer, and then resuspended in 50 µL Wash Buffer containing secondary antibody at a 1:50 dilution. Samples were incubated at 4°C for 1 hr, washed twice with 100 µL of Wash Buffer, and then resuspended in 50 µL 300-Wash Buffer with 1:50 diltuion of pA-Tn5. Samples were incubated at 4°C for 1 hr, washed 2X with 100 µL of 300-Wash Buffer, and then resuspended in 50 µL of Tagmentation Solution (300-Wash Buffer with 10 mM MgCl_2_). Samples were incubated at 37°C in a thermocycler with heated lid for 1 hr to allow the tagmentation reaction to go to completion. Samples were washed with 10 mM TAPS to remove any residual salt, and then resuspended in 10 mM TAPS pH8.5 containing 1X DAPI and 1X secondary diluent reagent (Takara Cat# 640196) at a concentration of 400 nuclei/µL. 80 µL of cell suspension was loaded into 8 wells of the 384 cell plate, together with 25 µL of the fiducial reagent (Takara Cat# 640196) according to the manufacturer’s instructions. Sample suspension (35 nL) was dispensed on the ICELL8 into the nanowells of a 350v Chip (Takara Cat# 640019). The 350v Chip was dried and sealed, and cells were centrifuged at 1200xg for 3 min. The Chip was then imaged to identify wells containing a single nuclei and a filter file was prepared. During image processing, 35 nL of 0.19% SDS in TAPS was added to all nanowells on the ICELL8 using an unfilitered dispense. The Chip was then dried, sealed and centrifuged at 1200xg for 3 min and then heated at 58°C in a thermocycler with heated lid for 1 hr to release the pA-Tn5 and prepare the tagmented chromatin for PCR. Before opening, the Chip was centrifuged at 1200xg, and 35 nL of 2.5% Triton-X100 neutralization solution was added to all wells containing a single nuclei via a filtered dispense on the ICELL8. The Chip was then dried and 35 nL of i5 indices was added via a filtered dispense. The Chip was then dried and 35 nL of i7 indices was added via a filtered dispense. The Chip was then dried, sealed and centrifuged at 1200xg for 3 min. Then 100 nL of KAPA PCR mix (2.775 X HiFi Buffer, 0.85 mM dNTPs, 0.05 U KAPA HiFi polymerase / µL)(Roche Cat# 07958846001) was added to all wells containing a single nucleus via two 50 nL filtered dispenses. The Chip was centrifuged at 1200xg for 3 min, sealed and placed in a thermocycler for PCR amplification using the following conditions: 1 cycle 58 ^0^C 5 min; 1 cycle 72 ^0^C 10 min; 1 cycle of 98 ^0^C 45 sec; 15 cycles of 98 ^0^C 15 sec, 60 ^0^C 15 sec, 72 ^0^C 10 sec; 1 cycle 72 ^0^C 2 min. The Chip was then centrifuged at 1200xg for 3 min into a collection tube (Takara Cat# 640048). To remove residual PCR primers and detergent, the sample was then cleaned up using two rounds of SPRISelect Ampure bead cleanup at a 1.3 : 1 v/v ratio of beads to sample. Samples were resuspended in 30 uL of 10 mM Tris-HCL pH 8.0, quantified on a Tapestation bioanalyzer instrument, and pooled with bulk samples for sequencing.

### DNA sequencing and Data processing

The size distributions and molar concentration of libraries were determined using an Agilent 4200 TapeStation. Up to 48 barcoded CUT&RUN libraries or 96 barcoded CUT&Tag libraries were pooled at approximately equimolar concentration for sequencing. Paired-end 25 × 25 bp sequencing on the Illumina HiSeq 2500 platform was performed by the Fred Hutchinson Cancer Research Center Genomics Shared Resources. This yielded 5-10 million reads per antibody. Single-cell CUT&Tag libararies were prepared using unique i5 and i7 barcodes and pooled with bulk samples for sequencing. For 500-100 cells 20 million reads was sufficient to obtain an average of approximately 80% saturation of the estimated library size for each single cell. Paired-end reads were aligned using Bowtie2 version 2.3.4.3 to UCSC HG19 with options: --end-to-end --very-sensitive --no-mixed -- no-discordant -q --phred33 -I 10 -X 700. Peaks were called from using SEACR (35) after combining replicates. We used custom scripts (https://github.com/FredHutch/SEACR/) to merge bulk histone modification-specific peak sets, map fragments to merged peak sets, and generate Principal Component Analyses (PCA) and t-Stochastic Neighbor Embeddings (t-SNE). All PCA was implemented using the prcomp() function in R (https://www.r-project.org/). t-SNE was implemented using the Rtsne() function in the Rtsne library. We used all principal components explaning greater than 1% of variance as input to Rtsne, and perplexity was set to the nearest integer to the square root of the number of rows in the input matrix. Bivalent gene classifications (H3K4me3-specific, H3K27me3-specific, and bivalent) for each cell type were determined by quantifying the number of reads mapping in a 2kb window around the TSS for every gene, and using a two-component Gaussian Mixture Model as implemented using the normalmixEM() function from the “mixtools” library in R to distinguish “enriched” and “non-enriched” sets of genes for each histone mark. Bivalent genes were designated as residing in the enriched gaussian component for both H3K4me3 and H3K27me3 in the cell type in question.

### Identifying *KMT2A*r oncoprotein targets

To identify unique *KMT2A*r targets, we first generated merged set of SEACR peaks originating from either N-terminal or C-terminal KMT2A antibody-targeted CUT&RUN in each cell type assayed. We quantified the number of fragments mapping to each peak i from each dataset j, and summed reads mapped from the two antibodies targeting the same KMT2A terminus in the same dataset to yield N-terminal (n_ij_) and C-terminal (c_ij_) fragments mapped in each peak, existing in cell type sets Nj and Cj, respectively. We calculated the cell type-specific “N over C ratio” (NCR) for each peak as follows:

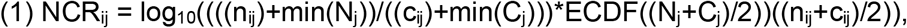

where min(x) = minimum value of x across the peak set; and ECDF(y)(x) = Empirical Cumulative Distribution Function of set y evaluated at x, as implemented in R using the ecdf() function. As illustrated in equation (1), ECDF was used to shrink NCR values towards zero in inverse proportion with the mean n_ij_+c_ij_ signal observed in the peak. *KMT2A*r identity was evaluated by fitting a two-component Gaussian Mixture Model to all NCR_j_ and asserting as True any NCR_ij_ that were greater than the NCR value greater than the mean of NCRj at which the two fitted Gaussian distributions intersect. As a second filter, the above Gaussian Mixture Modeling approach was repeated using peak length as an input, and peak_ij_ was considered a *KMT2Ar* oncoprotein-specific target only when both NCR and peak length met the cutoff described above. Gaussian Mixture Modeling was implemented in R using the normalMixEM() function from the “mixtools” library. For all peaks assigned as *KMT2A*r in any cell type, NCR scores were hierarchically clustered using the hclust() function in R on a euclidean distance matrix generated by the dist() function.

### t-SNE embedding of the active and repressed chromatin regions

For histone modification data, peaks were called from merged replicate datasets using SEACR^35^, and peak sets were merged for each modification across all cell types. We generated matrices of raw read counts mapping in each cell type (columns) to merged peaks (rows) for each modification, and we filtered out instances were counts were lower than any count value whose evaluated Empirical Cumulative Distribution Function was more than 5% diverged from the predicted ECDF value based on a lognormal fit of the data distribution, using the fitdistr() function from the MASS library with “densfun” set to “lognormal”. We then log_10_-transformed the results and rescaled columns to z-scores. Principal component analysis (PCA) was performed on the resulting transformed matrices using the prcomp() function in R. For t-SNE analysis, all principal components contributing greater than 1% variance were used as input to the Rtsne() function from the Rtsne library, with perplexity set as the nearest integer to the square root of the number of peaks, and check_duplicates set as FALSE. We used the resulting two-dimensional t-SNE values as input to the densityClust() function from the densityClust library, and used that output in the findClusters() function, with rho and delta values set to the 95^th^ percentile of all rho and delta values output from densityClust(), respectively. To generate cluster average heatmaps, scaled count values were averaged by cluster and the resulting matrix was used as input to the heatmap.2() function from the gplots library. PCA and t-SNE plots were generated using the ggplot2 library (https://ggplot2.tidyverse.org/).

### UMAP embedding of single cells

Single-cells that did not meet a minimum numbers of reads (n=300) or fell below the normal distribution of FRiPs defined by aggregate data were removed. Then, a single-cell count matrix of N features, defined by 5kb windows tiled across the genome, by M cells was generated. These matrices were then binarized and normalized via latent semantic indexing (LSI)^41^. The normalized count-matrix was reduced from N dimensions to two dimensions using UMAP and plotted. We generated imputed gene scores using MAGIC^42^ for subsequent analysis. Normalized dispersion was calculated from these gene scores using SCANPY^50^.

### Preparation of Figure Panels

All heat maps were generated using DeepTools^51^. t-SNE plots colored by maximum signal from immunophenotype class were generated using ggplot2. All of the data were analyzed using either bash, python (https://github.com/python), or R. The following packages were used in python: Matplotlib, NumPy, Pandas, Scipy, and Seaborn.

