## Supplementary Figures for "Automated CUT&Tag profiling of chromatin heterogeneity in mixed-lineage leukemia"

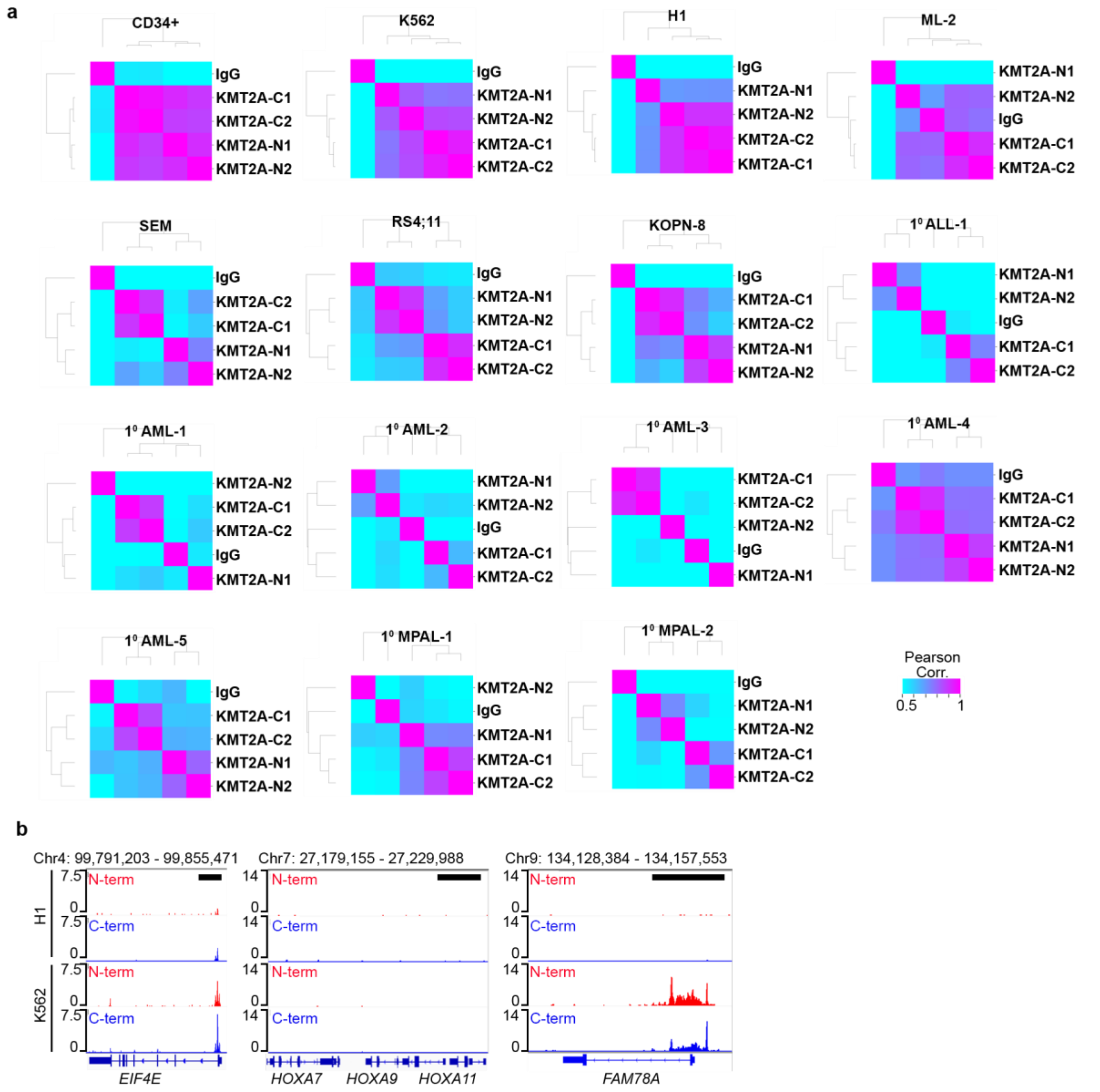


**Supplementary Figure 1: KMT2A N-terminus and C-terminus specific antibodies for AutoCUT&RUN chromatin profiling. a**, Pearson correlation matrices between KMT2A N-terminus and C-terminus specific antibodies over the KMT2A merged peaks for each sample. In the control CD34+ progenitors, as well as K562 and H1 cells signals for the KMTA-N1 antibody (Millipore Cat #05-764), KMTA-N2 antibody (Cell Signaling Tech Cat #14689S), KMTA-C1 antibody (Millipore Cat #05-765) and KMTA-C2 antibody (Santa Cruz Cat #sc-374392), are all highly correlated, indicating that in these samples the N-terminal and C-terminal regions of wildtype KMT2A co-localize on chromatin. In the *KMT2A*r samples profiles the N-terminal antibodies show higher correlations with one another than they do with profiles using the C-terminal antibodies, indicating the KMT2A-fusion binding is detected by the N-terminal antibodies and uncoupled from the KMT2A wildtype protein mapped by the C-terminal antibodies. **b**, Genome browser tracks showing examples of focused signal over the TSS of genes targeted by wildtype KMT2A in both H1 and K562 cells *(EIF4E)*. Critical hematopoietic cell fate determinants (*e.g. HOXA9*) are bound by wild type KMT2A in CD34+ HSPCs and the oncofusion proteins in *KMT2A*r leukemia, but are not bound by wildtype KMT2A in H1 or K562 cells. A broad distribution of KMT2A signal is found across the gene bodies of limited collection of target genes (e.g. *FAM78A*) in K562 cells. Black scale bars = 10 kb.

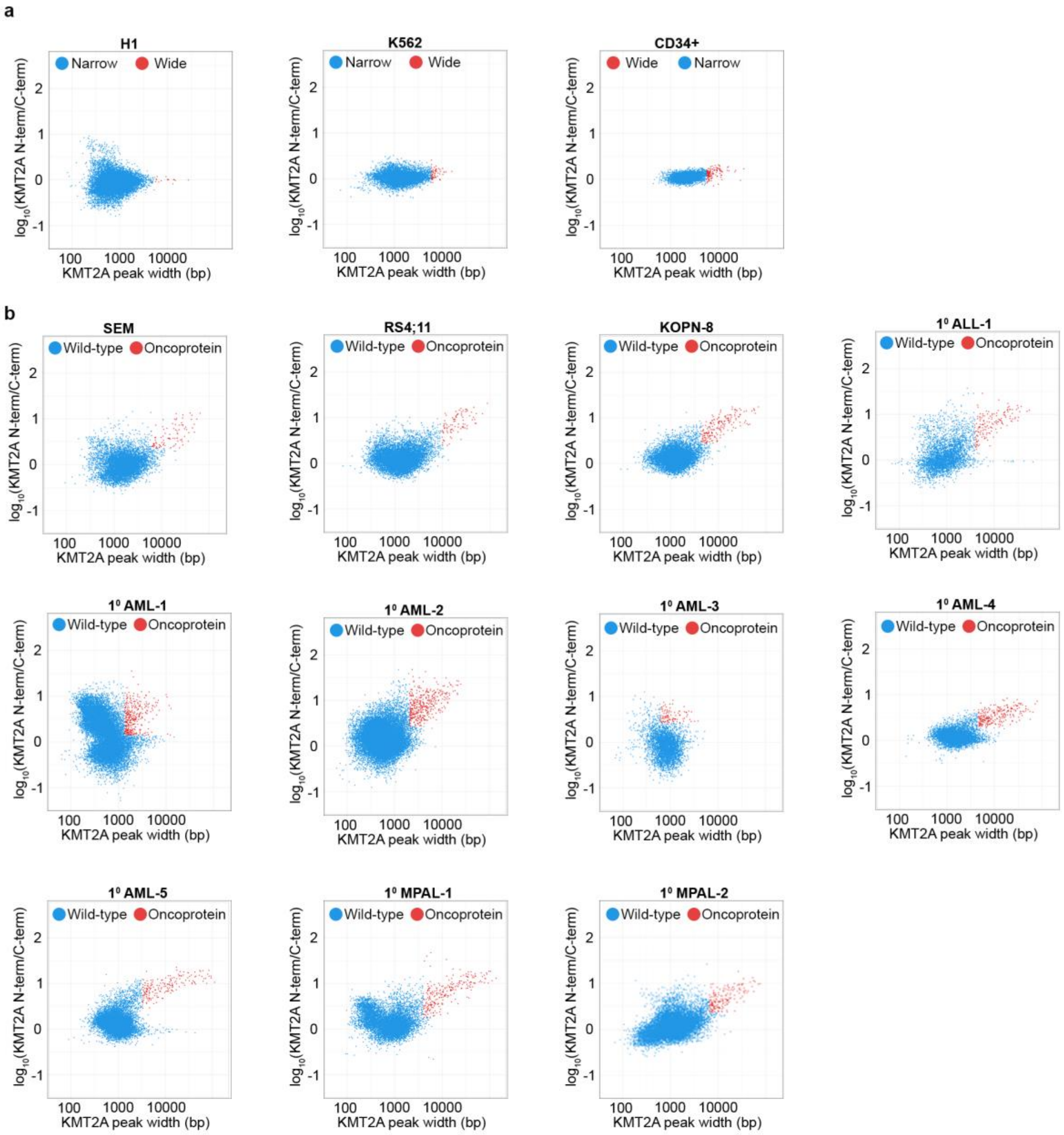


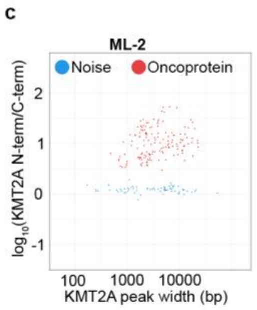


**Supplementary Figure 2: A uniform approach to identify sites bound by the KMT2A-fusion protein using KMT2A N- and C-terminal profiles a,** KMT2A scatter plots comparing the peak width and relative enrichment of the KMT2A N- versus C-terminus in control H1, K562, and CD34+. Wide peaks called by a two-component Gaussian mixture model of peak width are indicated in red. **b**, Same as (a) but for *KMT2A*r leukemia samples. Fusion oncoprotein targets we called using a two-component Gaussian mixture model of both peak width and N/C scores, which are indicated in red. **c**, Scatterplot of N/C scores called using a two-component Gaussian mixture model as in (a) for the KMT2A-AF6 containing ML-2 control cell line in which the wildtype *KMT2A* allele is deleted.

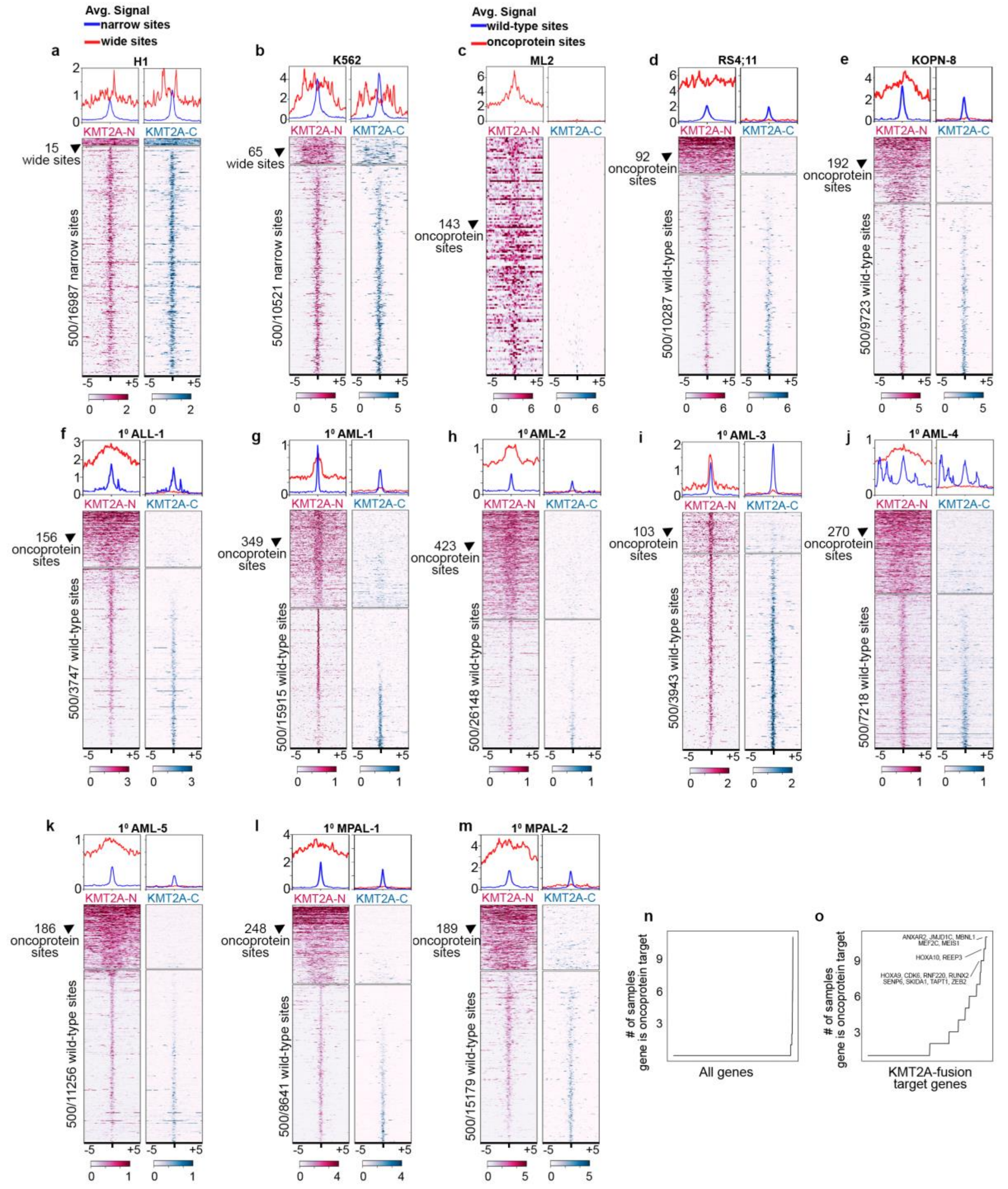

**Supplementary Figure 3: Comparison of fusion oncoprotein binding sites across all samples. a,** Heatmaps show the even distribution of the KMT2A N- and C-terminal profiles across the wide sites called in the control H1 cells (top). A random sample of 500 narrow peaks are shown for comparison (bottom). **b,** Same as (a) but in the control K562 cells. **c,** In the ML-2 cell line only the N-terminal profiles show appreciable signal over the fusion oncoprotein bound sites, which is consistent with the lack of wildtype KMT2A in these samples. **d**, In RS;411 cells the fusion oncoprotein bound sites (top) show a broad enrichment for the KMT2A N-terminal profiles and are severaly depleted for the C-terminal profiles. A random sample of 500 wildtype sites are shown for comparison (bottom) and show a similar overall enrichment of both the N- and C-terminal KMT2A profiles. **e-m,** Same as (d) but for the indicated *KMT2A*r samples.

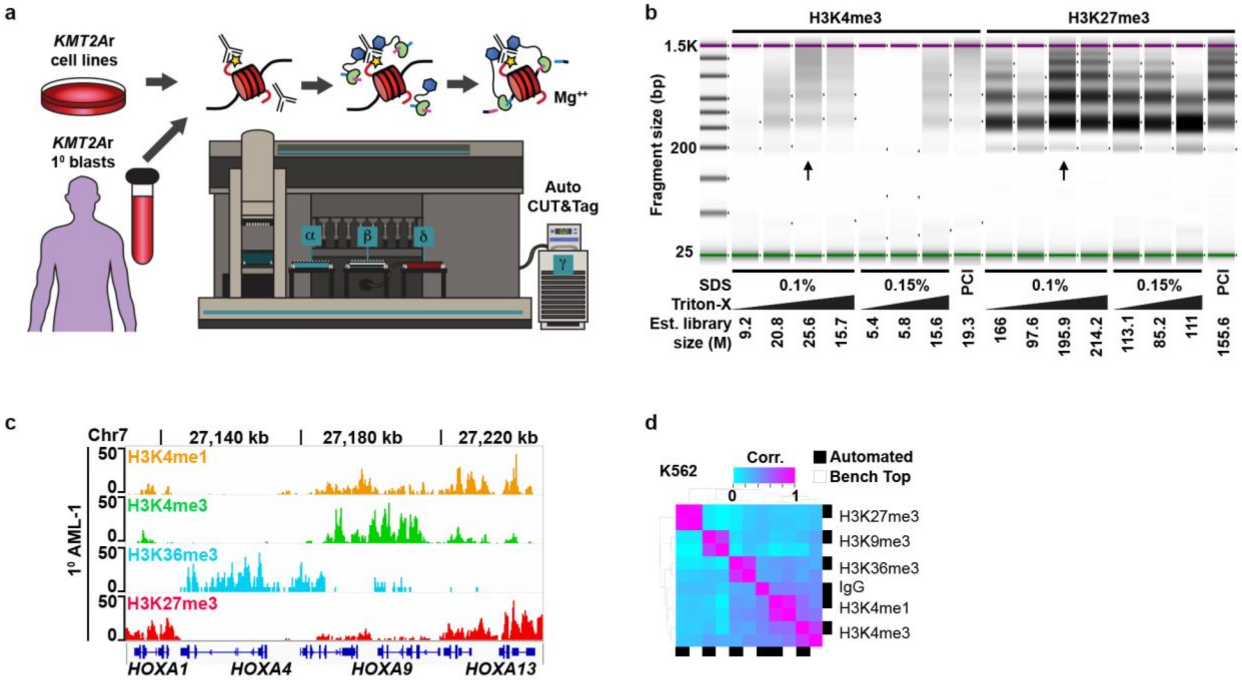


**Supplementary Figure 4: Adaptation of CUT&Tag for full automation. a,** Con-A bound nuclei are incubated with the primary antibody of interest and arrayed for AutoCUT&Tag profiling on a liquid handling robot equipped for high volume magnetic separation ($\alpha$), low volume magnetic separation ($\beta$) and temperature control ($\delta$, $\gamma$). This method pepares up to 96 sequencing-ready samples in a single day that can all be pooled on a single HiSeq two-lane or comparable flow cell for sequencing. The automated protocol uses a low concentration of SDS to displace bound Tn5 from tagmented DNA, and Triton-X100 to quench the detergent for PCR. **b,** A titration experiment showing optimization of the DNA release and quenching conditions for AutoCUT&Tag. Varying amounts of SDS and Triton-X100 were tested for library yield. Arrows indicate the optimum condition. **c,** Segmentation of a representative region of the human genome in the $1^0$ AML-1 patient sample by the histone modifications used in this study. **d,** Pearson correlation matrix of reproducibility between benchtop and automated CUT&Tag profiling methods on K562 fixed nuclei with 5 antibodies to histone modifications as well as the IgG negative control antibody.

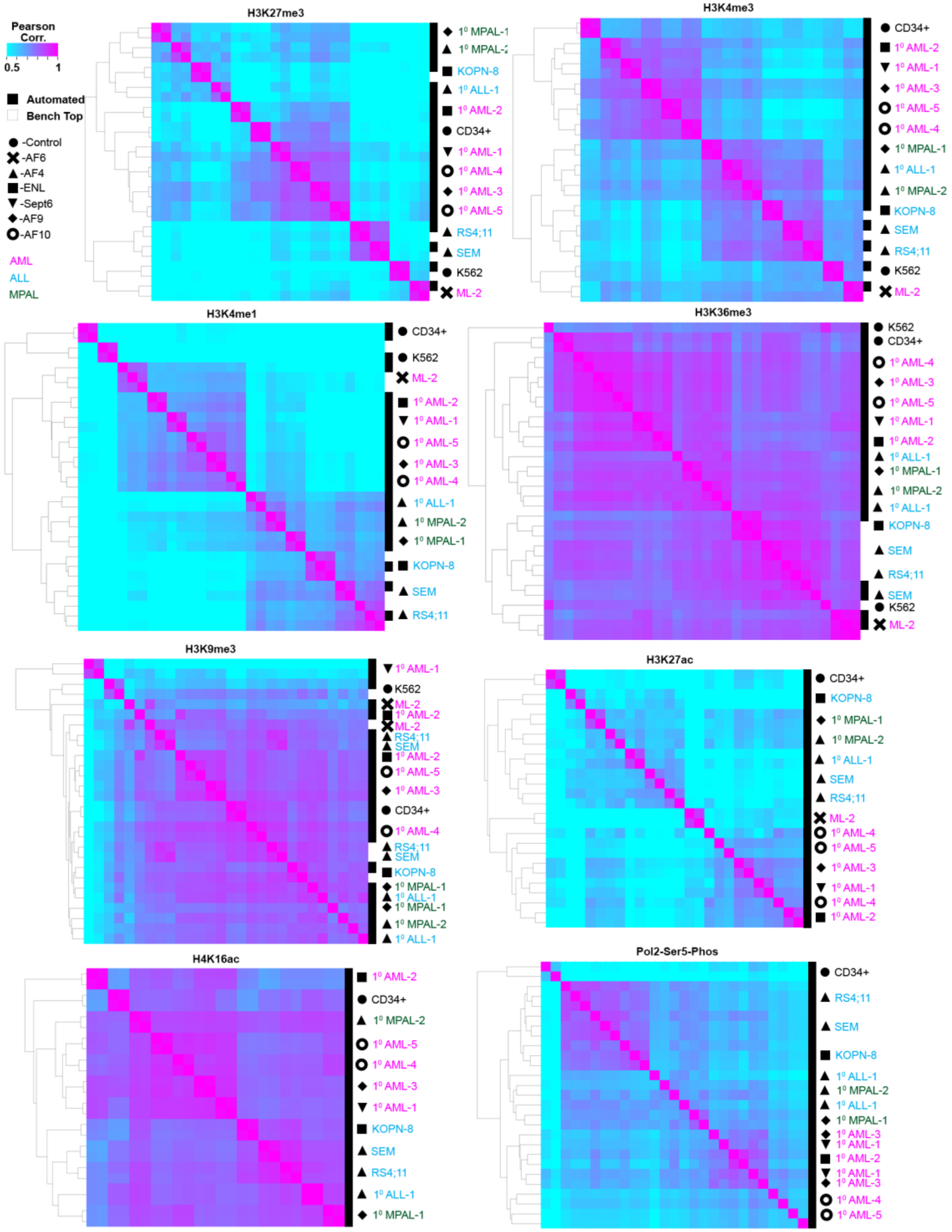


**Supplementary Figure 5: AutoCUT&Tag is highly reproducible for profiling primary patient leukemia. a,** Pearson correlation matrix of reproducibility between benchtop and automated CUT&Tag profiling methods on fixed nuclei from all 12 *KMT2A*r leukemias as well as the CD34+ progenitor and K562 control samples using antibodies against H3K27me3, H3K4me3, H3K4me1, H3K36me3, H3K9me3, H3K27ac, H4K16ac, and RNAP2S5p. H3K27me3, H3K4me3, H3K4me1, H3K27ac, and RNAP2S5p show the highest variation between samples, and most reliably cluster sample replicates together.

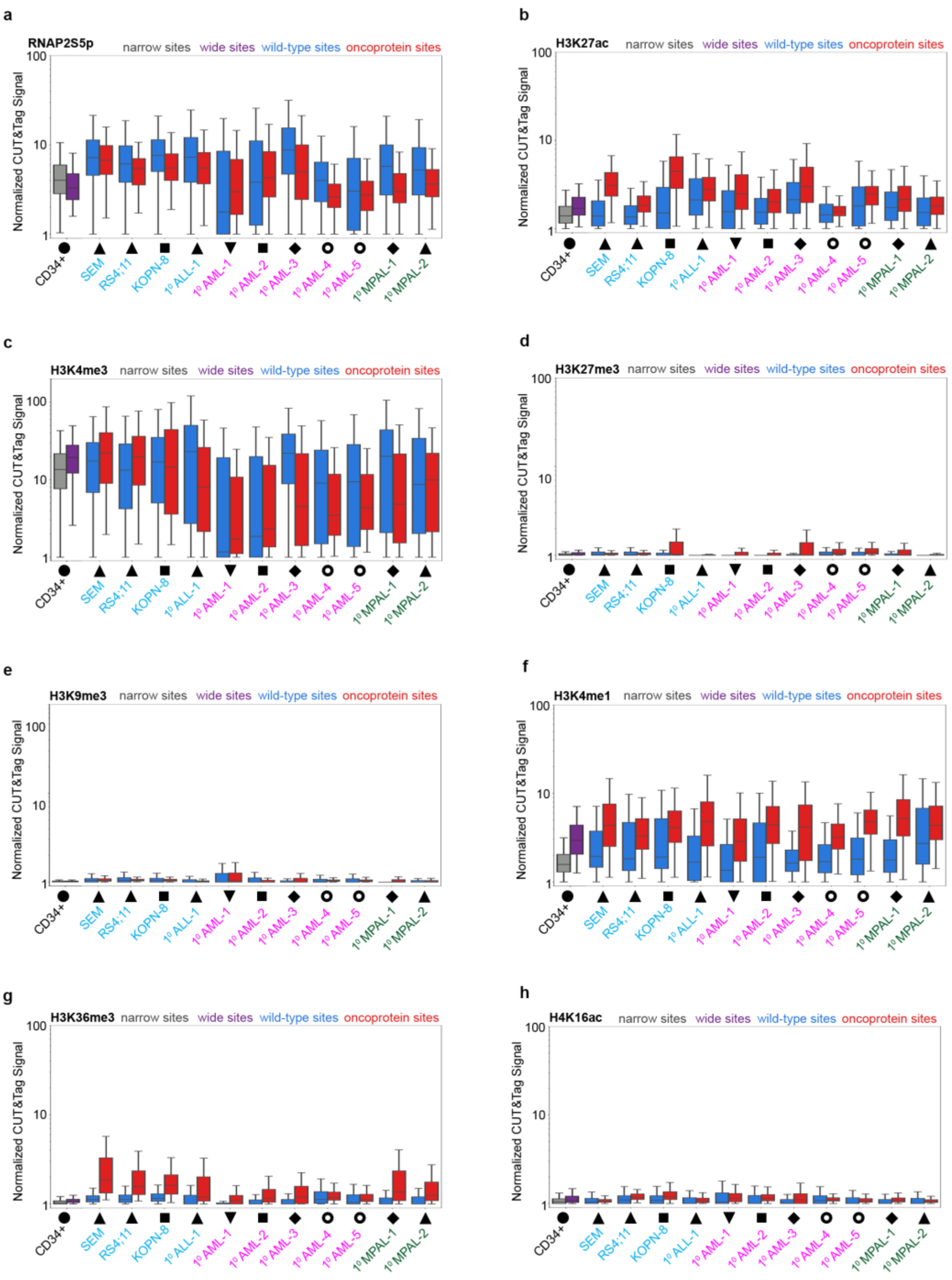


**Supplementary Figure 6: Chromatin features of KMT2A fusion-protein binding sites. a**, Quantification of the RNAP2Sp signal across the wildtype and oncoprotein KMT2A target sites in all samples. Signal is normalized by total coverage, as well as peak width. **b**, Same as (a) but for H3K27ac. H3K27ac is significantly enriched at the fusion oncoprotein binding sites in the SEM (median wildtype sites = 1.40, median oncoprotein sites =3.11, p-value = 7.71e-10), KOPN-8 (median wildtype sites =1.51, median oncoprotein sites =4.47, p-value = 2.41e-11), $1^{0}$AML-1 (median wildtype sites =1.55, median oncoprotein sites =2.49, p-value = 8.63e-13), $1^{0}$AML-2 (median wildtype sites =1.53, median oncoprotein sites =2.00, p-value = 2.76e-8), $1^{0}$AML-3 (median wildtype sites =2.12, median oncoprotein sites =2.99, p-value = 6.00e-8), and $1^{0}$AML-5 (median wildtype sites =1.81, median oncoprotein sites =2.45, p-value = 7.71e-10) *KMT2A*r leukemia samples as compared to the sample mached wildtype KMT2A bound sites. **Supplementary Figure 6 Legend continued on next page.**

**Supplementary Figure 6 Legend continued.** **c**, Same as (a) but for H3K4me3. H3K4me3 is significantly enriched at the fusion oncoprotein binding sites in SEM (median wildtype sites = 17.45, median oncoprotein sites =22.12, p-value = 4.15e-5), RS4;11 (median wildtype sites = 13.25, median oncoprotein sites =19.67, p-value = 0.042), $1^0$AML-1 (median wildtype sites = 1.17, median oncoprotein sites =1.72, p-value = 0.035), $1^0$AML-2 (median wildtype sites = 1.86, median oncoprotein sites =2.30, p-value = 0.0051) and $1^0$MAPL-2 (median wildtype sites = 8.67, median oncoprotein sites =9.95, p-value = 0.00028) and significantly depeleted in $1^0$ALL-1 (median wildtype sites = 22.94, median oncoprotein sites =8.09, p-value = 3.49e-9), $1^0$AML-3 (median wildtype sites = 22.00, median oncoprotein sites =4.51, p-value = 0.0019), $1^0$AML-4 (median wildtype sites = 9.04, median oncoprotein sites =3.48, p-value = 3.09e-11), $1^0$AML-5 (median wildtype sites = 9.38, median oncoprotein sites =4.34, p-value = 1.16e-10) and $1^0$MPAL-1 (median wildtype sites = 20.02, median oncoprotein sites =4.80, p-value = 2.22e-13). **d**, Same as (a) but for H3K27me3. **e**, Same as (a) but for H3K9me3. **f**, Same as (a) but for H3K4me1. **g**, Same as (a) but for H3K36me3. **h**, Same as (a) but for H4K16ac. Statistics are included in Supplementary Table 3.

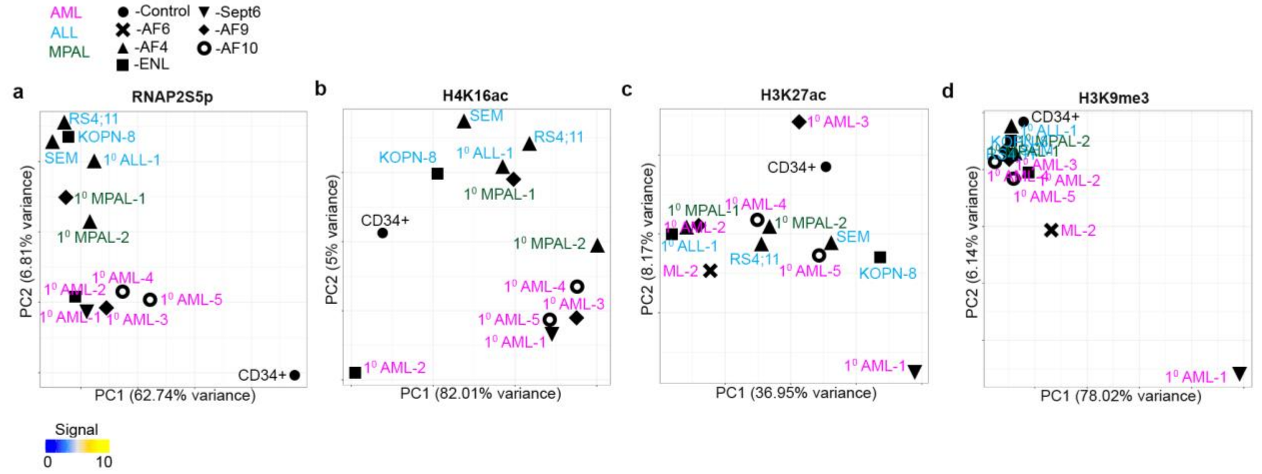


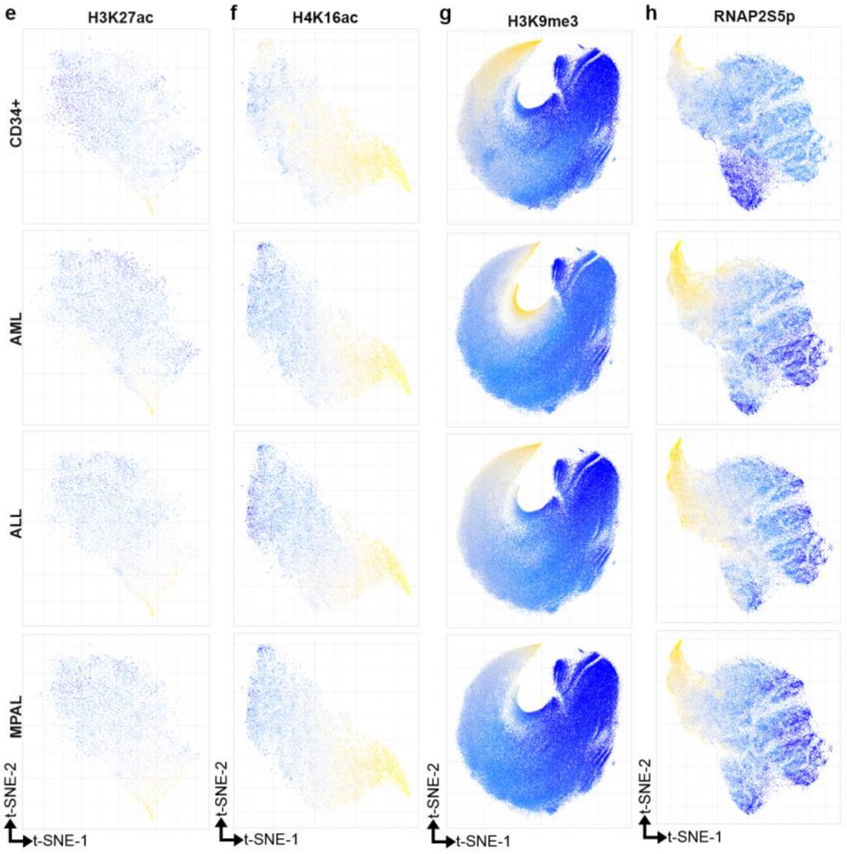


**Supplementary Figure 7: Grouping *KMT2A*r samples and regulatory regions according to the sample specific AutoCUT&Tag chromatin profiles.**

**a,** PCA of genome-wide RNAP2S5p signal separates samples by leukemia subtype. **b**, PCA of genome-wide H4K16ac signal separates samples by leukemia subtype. **c**, Samples organized by PCA of H3K27ac are not separated by leukemia subtype. **d**, PCA of genome-wide H3K9me3 signal only distinguishes the $1^0$ AML-1 KMT2A-Sept6 containing sample from the rest. **e-g**, t-SNE projections of H3K27ac H4K16ac and H3K9me3 samples do not organize the enriched elements based on enrichment in specific lineages. **h**, t-SNE projection of the RNAP2S5p bound regions identifies groups of elements that show CD34+, AML, ALL and MPAL specific enrichment of RNAP2S5p.

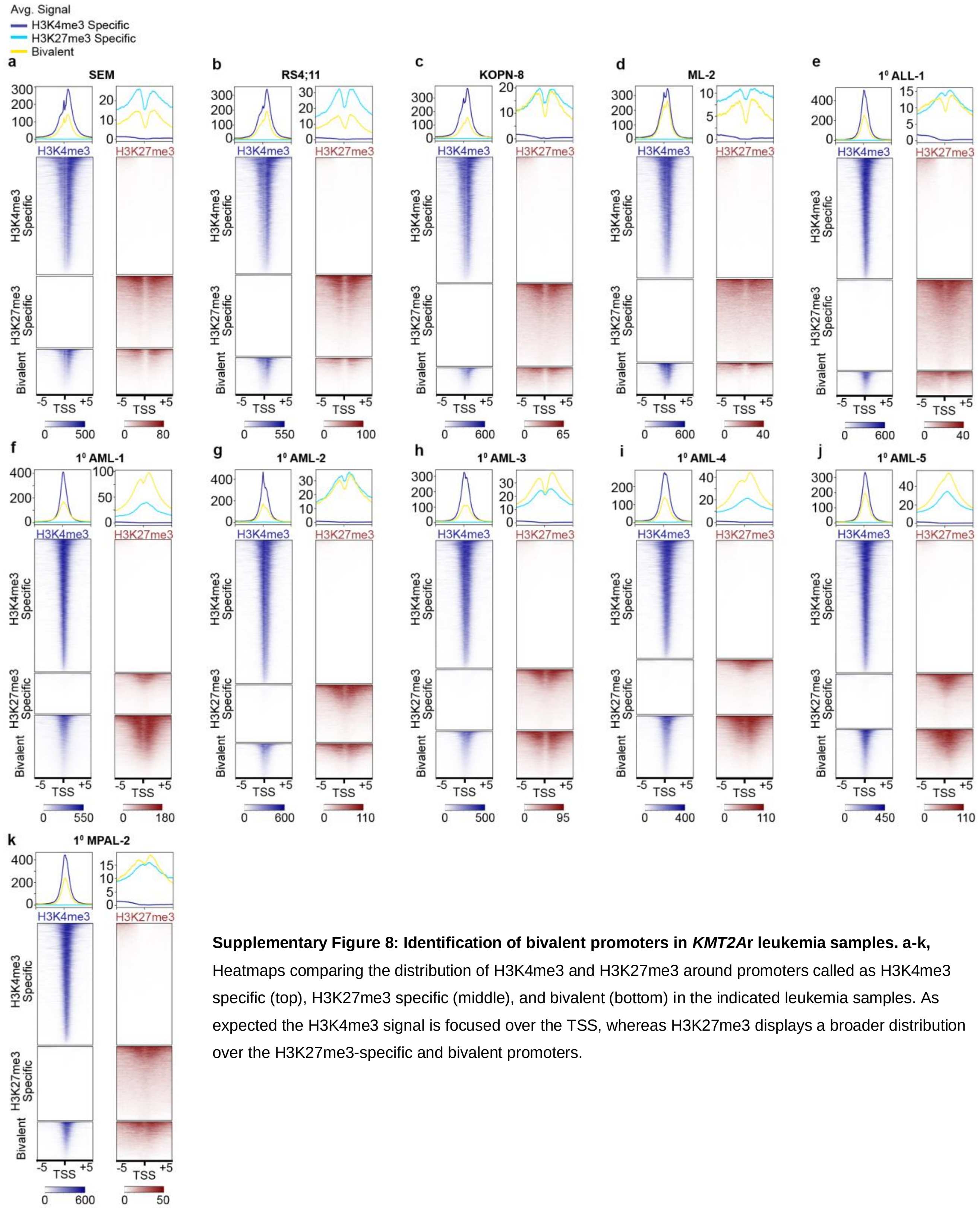


**Supplementary Figure 8: Identification of bivalent promoters in *KMT2A*r leukemia samples. a-k,** Heatmaps comparing the distribution of H3K4me3 and H3K27me3 around promoters called as H3K4me3 specific (top), H3K27me3 specific (middle), and bivalent (bottom) in the indicated leukemia samples. As expected the H3K4me3 signal is focused over the TSS, whereas H3K27me3 displays a broader distribution over the H3K27me3-specific and bivalent promoters.

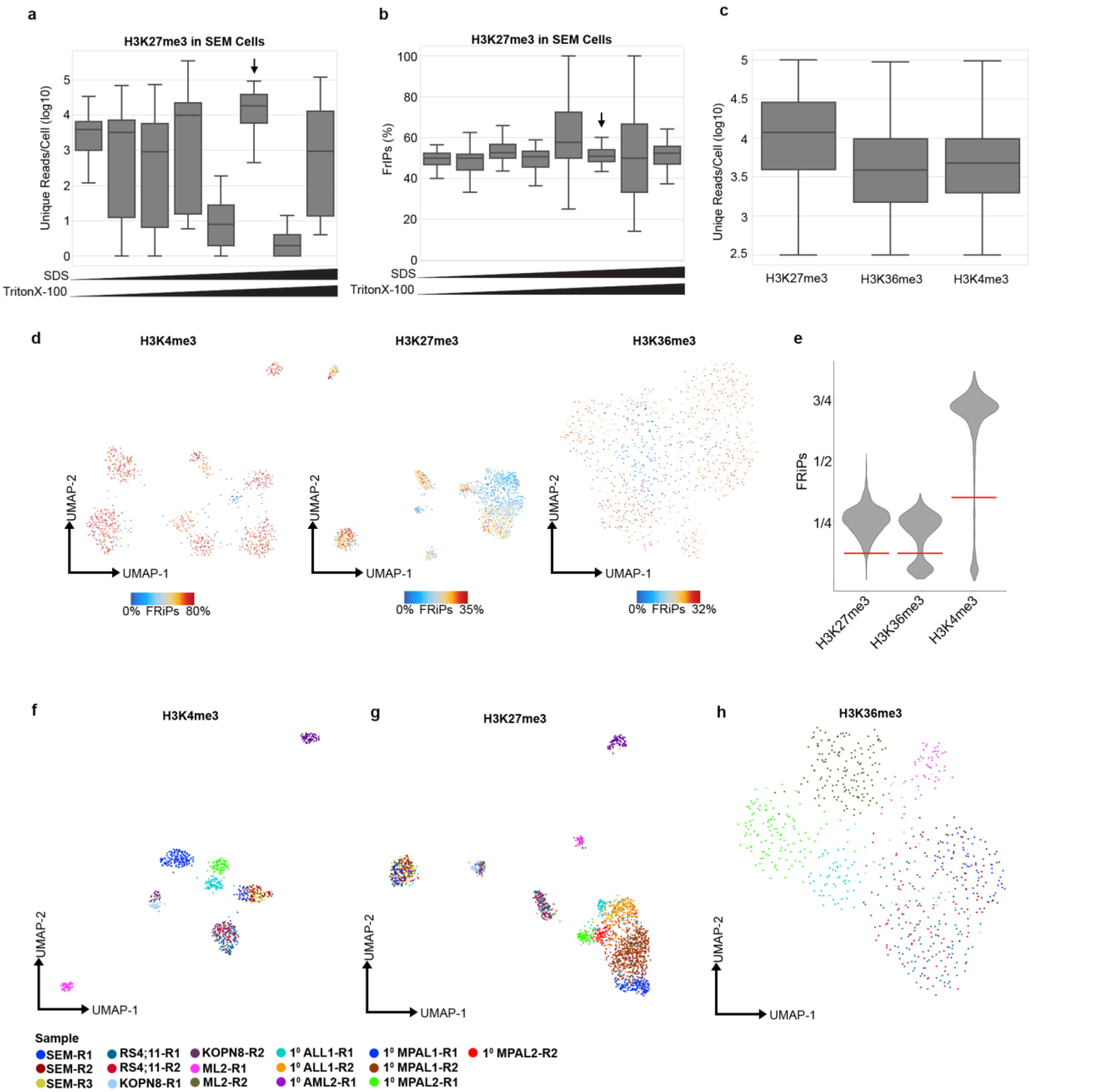


**Supplementary Figure 9: Optimization of CUT&Tag-Direct for single cell applications on the ICELL8. a,** Titrating the concentration of SDS and Triton-X in the nanowell increases the library yield for individual cells, and identifies optimum conditions for Tn5 release and PCR enrichment (Arrow). **b,** The Fraction of Reads in Peaks (FRiPs) varies across SDS and Triton-X titration conditions. Arrow indicates the optimum conditions for Tn5 release and PCR enrichment. **c**, Boxplots showing the distributions of unique reads per cell across all of the cells profiled for H3K27me3, H3K36me3 and H3K4me3. **d,** UMAP projection of single cells profiled with H3K4me3, H3K27me3 or H3K36me3 colored according to the FRiP scores of each individual cell using peaks called on the aggregate data of all cells profiled for a given mark. Cells with low FRiP scores tend to fall in between clusters and were removed as a quality control. **e**, Violin plot showing the distribution of FRiP scores for all individual cells profiled using the indicated histone mark. The red lines indicate the cut-offs used for quality control for each mark. **f**, UMAP projection of single cells profiled for H3K4me3 and colored according to batch (R1, R2, R3). Cells from the same sample processed on different days cluster together in UMAP space indicating that batch effects have a minimal impact on the separation of cells according to the H3K4me3 profiles. **g**, Same as (f) for H3K27me3. **h**, Same as (f) for H3K36me3.

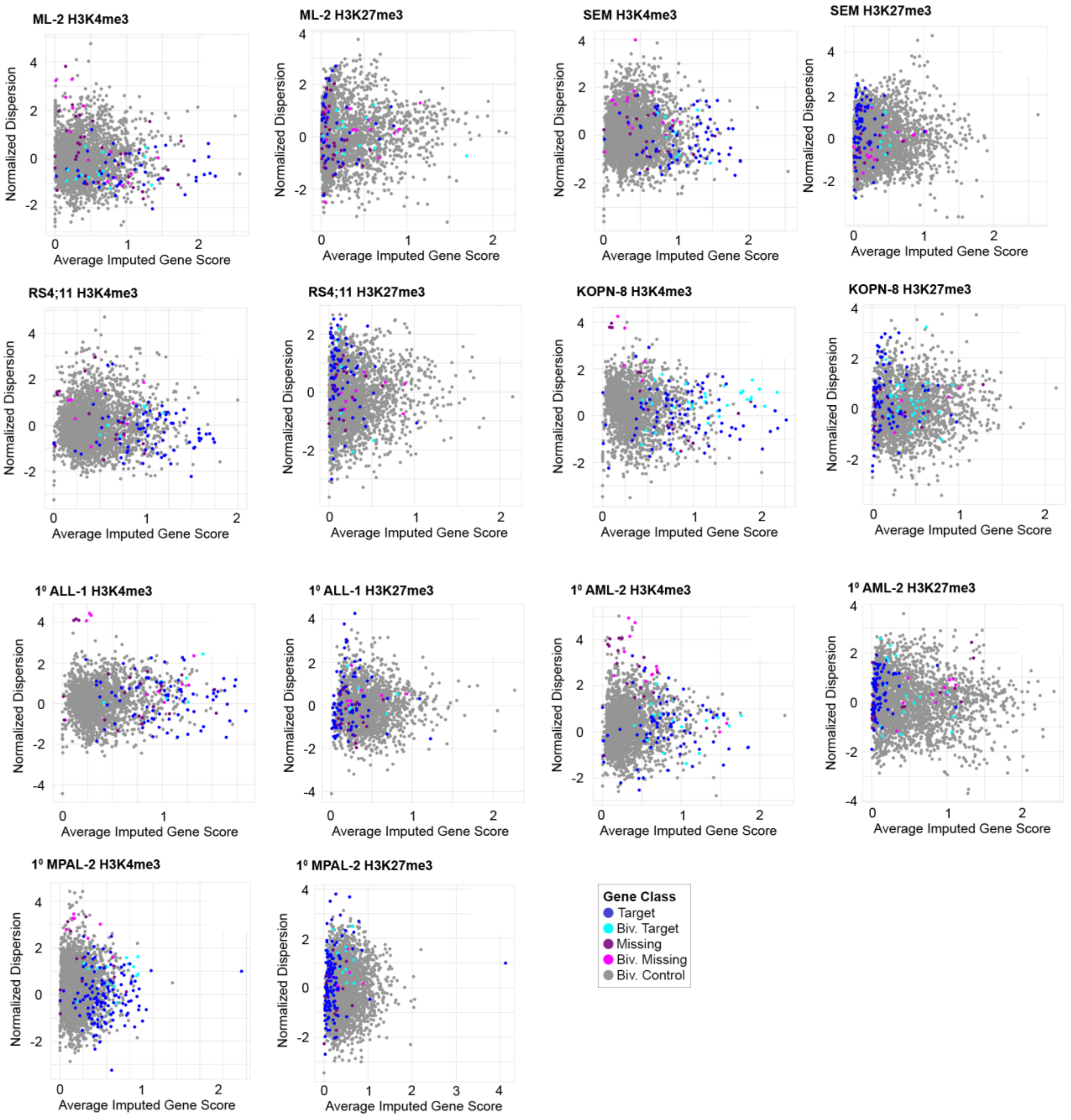


**Supplementary Figure 10: Heterogeneous enrichment of H3K4me3 and H3K27me3 at bivalent fusion oncoprotein targets in *KMT2A*r leukemia samples.** Scatterplots comparing the average imputed H3K4me3 scores or H3K27me3 scores and normalized dispersion of genes grouped according to the KMT2A-fusion binding status (Target, Missing Target, and unbound Control) and promoter bivalency status (Biv.) within the indicated samples.

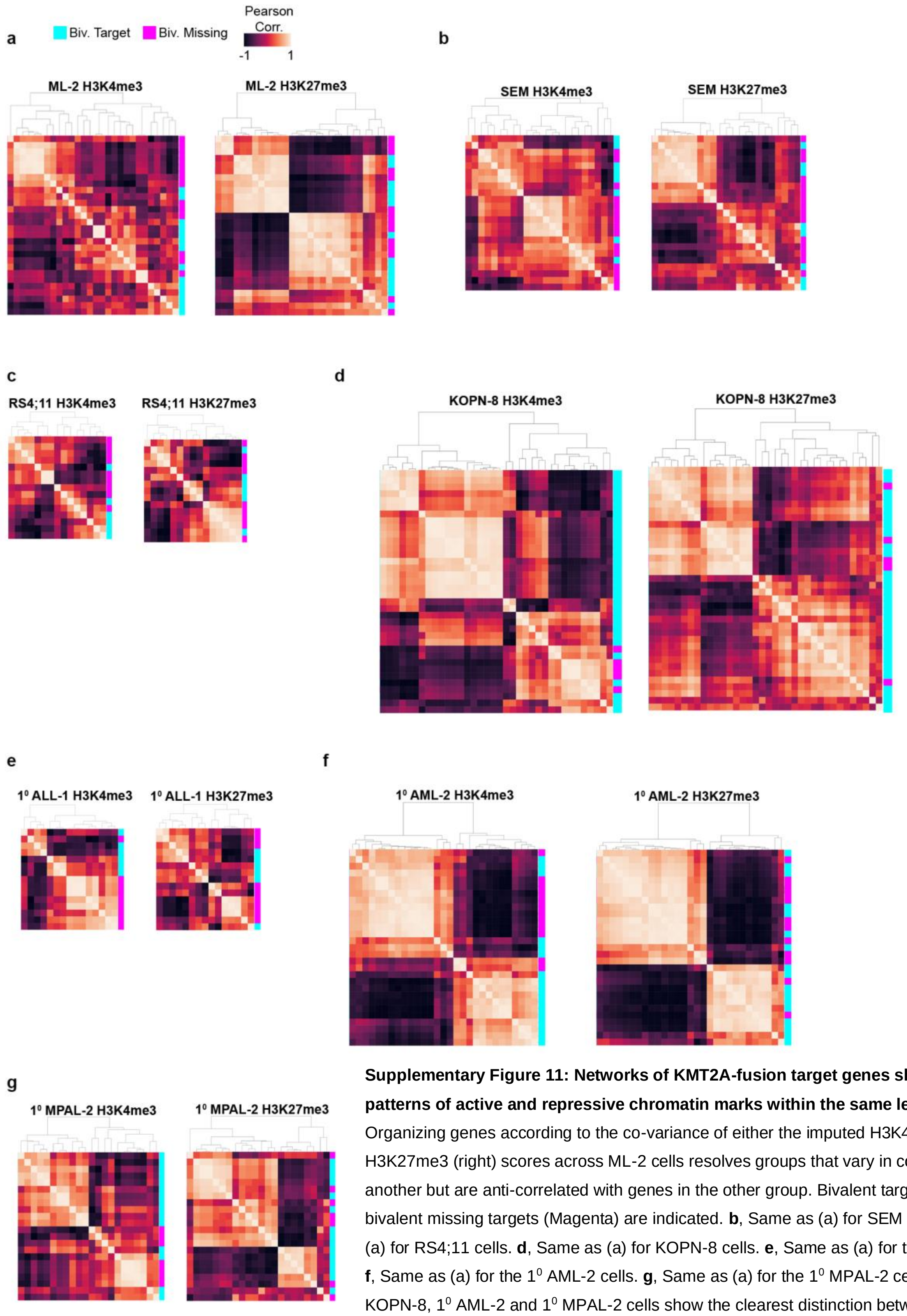


**Supplementary Figure 11: Networks of KMT2A-fusion target genes show divergent patterns of active and repressive chromatin marks within the same leukemia. a**, Organizing genes according to the co-variance of either the imputed H3K4me3 (left) or H3K27me3 (right) scores across ML-2 cells resolves groups that vary in concert with one another but are anti-correlated with genes in the other group. Bivalent targets (Cyan) and bivalent missing targets (Magenta) are indicated. **b**, Same as (a) for SEM cells. **c**, Same as (a) for RS4;11 cells. **d**, Same as (a) for KOPN-8 cells. **e**, Same as (a) for the $1^0$ ALL-1 cells. **f**, Same as (a) for the $1^0$ AML-2 cells. **g**, Same as (a) for the $1^0$ MPAL-2 cells. The ML-2, KOPN-8, $1^0$ AML-2 and $1^0$ MPAL-2 cells show the clearest distinction between divergent groups of oncoprotein target genes.
